# m^6^A RNA methylation orchestrates transcriptional dormancy during developmental pausing

**DOI:** 10.1101/2023.01.30.526234

**Authors:** Evelyne Collignon, Brandon Cho, Julie Fothergill-Robinson, Giacomo Furlan, Robert L. Ross, Patrick A. Limbach, Miguel Ramalho-Santos

## Abstract

Embryos across metazoan lineages can enter reversible states of developmental pausing, or diapause, in response to adverse environmental conditions. The molecular mechanisms that underlie this remarkable dormant state remain largely unknown. Here we show that m^6^A RNA methylation by Mettl3 is required for developmental pausing in mice by maintaining dormancy of paused embryonic stem cells and blastocysts. Mettl3 enforces transcriptional dormancy via two interconnected mechanisms: i) it promotes global mRNA destabilization and ii) suppresses global nascent transcription by specifically destabilizing the mRNA of the transcriptional amplifier and oncogene N-Myc, which we identify as a critical anti-pausing factor. Our findings reveal Mettl3 as a key orchestrator of the crosstalk between transcriptomic and epitranscriptomic regulation during pausing, with implications for dormancy in stem cells and cancer.

## Main Text

Development is often described as a sequential unfolding of genetic programs towards increased complexity, occurring with stereotypical timing. However, many animal species can pause early embryonic development to improve survival under adverse environmental conditions by entering a dormant state called embryonic diapause(*1, 2*). In mammals, pausing manifests as a delayed implantation of the blastocyst, the source of pluripotent embryonic stem cells (ESCs)(*3, 4*). This state of paused pluripotency can be induced in mouse blastocysts and ESCs by the inhibition of mTOR, a conserved growth-promoting kinase, and is characterized by a drastic decrease in proliferation and biosynthetic activity, including transcription(*5*). However, a detailed characterization of the mechanisms regulating transcription in this context is still missing.

To gain insights into the regulation of transcriptional dormancy during pausing, we performed a comprehensive screen for chemical modifications of RNA in paused ESCs. Mass spectrometry revealed significantly increased levels of N^6^-methyladenosine (m^6^A) in paused ESCs, relative to control condition (Fig. 1A, fig. S1A). This modification is deposited on nascent RNA by a methyltransferase complex, with Mettl3 as the catalytically active subunit(*6, 7*). m^6^A plays essential roles during post-implantation development via mRNA destabilization of key cell fate regulators(*8–10*). We hypothesized that the increase in m^6^A may indicate an unexpected role for this mark during paused pluripotency.

**Fig. 1.**
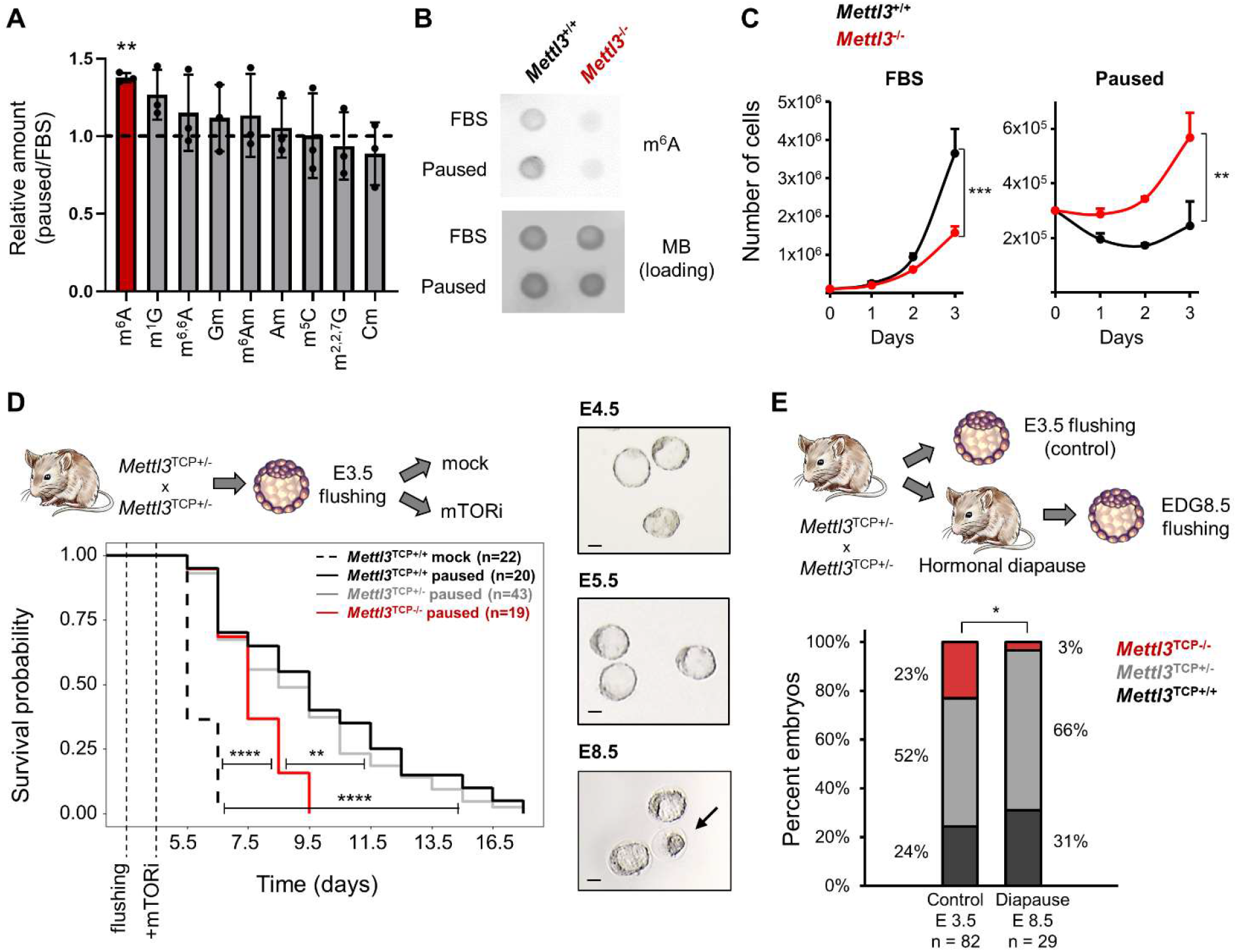
The m_6_A methyltransferase Mettl3 is essential for paused pluripotency. (**A**) Screening of RNA modifications by mass spectrometry in paused ESCs, normalized to FBS-grown ESCs. Data are mean ± SD (n=3). Ratio paired Student ’s *t*-tests \*\**P* < 0.01. (**B**) Dot blot showing that the increase of m^6^A in paused ESCs is abrogated in *Mettl3*^-/-^. (**C**) Growth curves showing that *Mettl3*^-/-^ ESCs fail to suppress proliferation in paused conditions. Data are mean ± SEM (n=3). Linear regression test \*\**P* < 0.01, \*\*\**P* < 0.001. (**D**) *Mettl3* loss leads to the premature death of mouse blastocysts cultured ex vivo in paused conditions. Log-rank test \*\**P* < 0.01, \*\*\*\**P* < 0.0001. (**E**) Quantification of recovered (live) embryos at E3.5 (control) and at Equivalent Days of Gestation (EDG) 8.5 following hormonal diapause, showing that *Mettl3*^TCP-/-^ embryos are impaired at undergoing hormonal diapause. χ2 test \**P* < 0.05.

Paused ESCs, induced by mTOR inhibition, are viable and pluripotent but proliferate very slowly(*5*). However, we found that paused *Mettl3*^-/-^ ESCs grow at a much faster rate than paused wildtype (*Mettl3*^+/+^) ESCs, suggesting defective suppression of proliferation (Fig. 1B-C, fig. S1B-F). Interestingly, this faster proliferation rate of *Mettl3*^-/-^ ESCs is observed only in the paused state and not in control FBS conditions (Fig. 1C, fig. S1D,F), suggesting a specific role in developmental pausing. As we reported previously, inhibition of mTOR prolongs survival of blastocysts ex vivo for 1-2 weeks and induces a paused state(*5*), a finding reproduced here with *Mettl3*^+/+^ embryos (Fig. 1D). By contrast, we found that *Mettl3*^-/-^ blastocysts are prematurely lost during ex vivo pausing (Fig. 1D and see model in fig. S1G-H). *Mettl3*^-/-^ embryos are also largely incompatible with hormonally induced diapause (Fig. 1E). Taken together, these findings reveal an essential role for Mettl3 in ESC and blastocyst pausing.

A central feature of paused pluripotency is a global reduction in transcriptional output, or hypotranscription, encompassing both coding and non-coding RNAs(*5*). In comparison with paused *Mettl3*^+/+^ cells, paused *Mettl3*^-/-^ ESCs display increased levels of both total and nascent RNA per cell (Fig. 2A-C, fig. S2A). Levels of nascent RNA are also elevated in paused *Mettl3*^-/-^ blastocysts, as compared with *Mettl3*^+/-^ or *Mettl3*^+/+^ embryos (Fig. 2D-E). We next performed cell number-normalized RNA-sequencing, which uses exogenous RNA spike-ins and allows for quantification of global shifts in transcriptional output(*11*) (fig. S2B, Table S1, and see Methods). In line with the global changes observed in RNA levels (Fig. 2A-C), paused *Mettl3*^-/-^ ESCs displayed a defective hypotranscriptional state (Fig. 2F, fig. S2C-E). Indeed, while 10,656 genes are downregulated in paused *Mettl3*^+/+^ cells, only 5,916 genes (i.e. 55.5%) are downregulated in paused *Mettl3*^-/-^ cells (fig. S2D). This suppression of hypotranscription is particularly evident for pathways and functional categories whose silencing is a feature of developmental pausing, such as translation, ribosome biogenesis, mTOR signaling, Myc targets and energy metabolism(*4, 5, 12–14*) (Fig. 2G-H, fig. S2F-G, Table S2). These results reveal that Mettl3 contributes to the global transcriptional dormancy observed during developmental pausing.

**Fig. 2.**
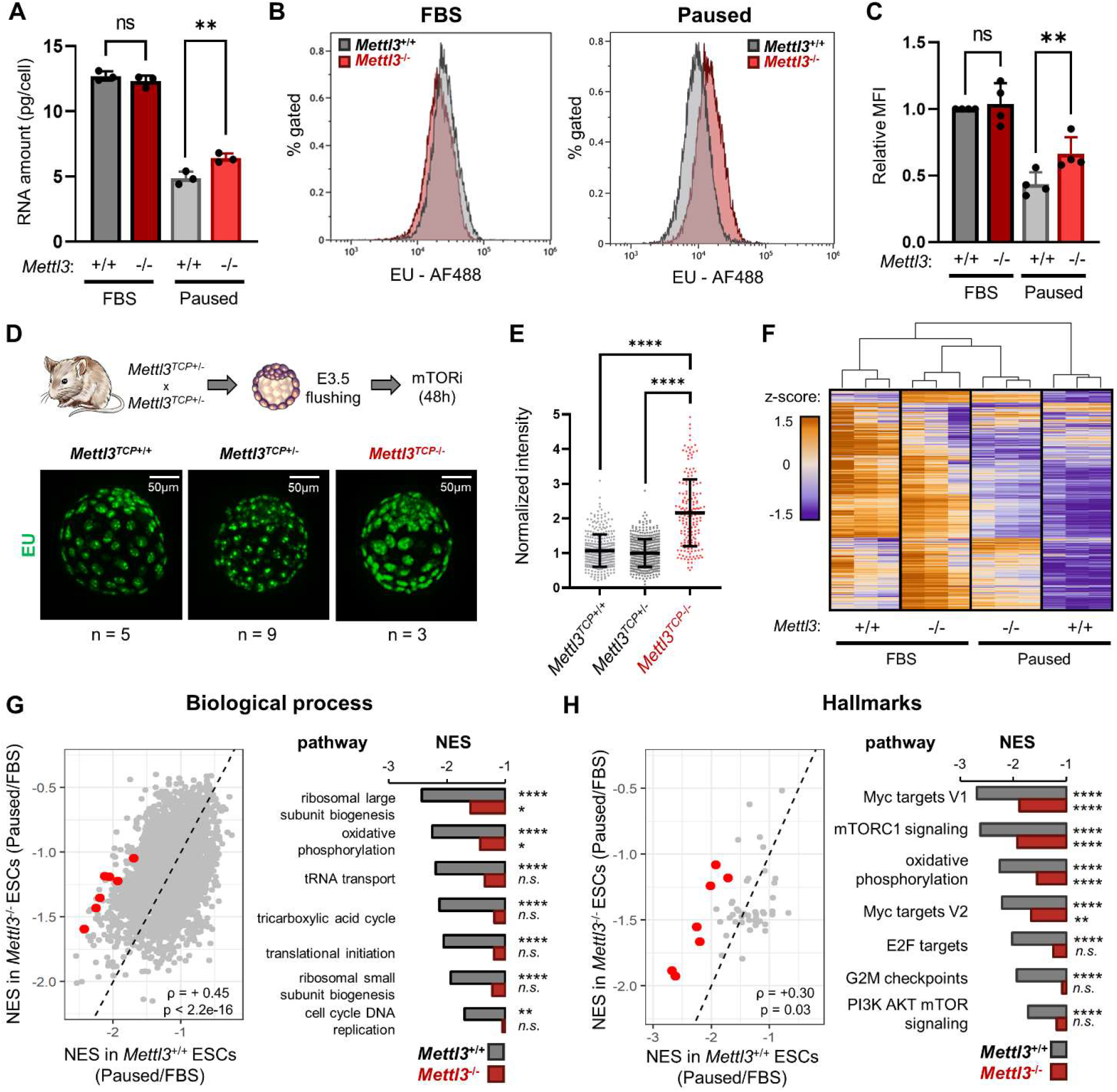
Mettl3 regulates hypotranscription in paused pluripotency. (**A**) Total RNA per cell in *Mettl3*^+/+^ and *Mettl3*^-/-^ ESCs, in FBS and paused conditions. Data are mean ± SD, (n=3). Paired Student ’s *t*-tests \*\**P* < 0.01. (**B**,**C**) Nascent transcription by EU incorporation, quantified by median fluorescence intensity (MFI) relative to *Mettl3*^+/+^ FBS in each experiment. Data are mean ± SD (n=4). Paired Student ’s *t*-tests \*\**P* < 0.01. (**D**,**E**) Immunofluorescence and nuclear signal quantification of EU incorporation in ex vivo paused blastocysts, showing increased nascent transcription in *Mettl3*^*TCP*-/-^. Data are mean ± SD. One-way ANOVA with Dunnett ’s multiple comparison test \*\*\*\**P* < 0.0001. (**F**) Heatmap of gene expression for all genes expressed in ESCs, showing defective hypotranscription in paused *Mettl3*^-/-^ ESCs. (**G**,**H**) Gene set enrichment analysis (GSEA) of gene expression changes in paused ESCs, using the “GO biological processes” (G) and “hallmarks” collections (H). Scatter plots of the normalized enrichment scores (NES), with Spearman correlation coefficient (ρ) with representative pathways showing defective hypotranscription in *Mettl3*^-/-^ (red dots) highlighted. pre-ranked gene set enrichment analysis with FDR correction \**P* < 0.05, \*\**P* < 0.01, \*\*\*\**P* < 0.0001.

To probe the mechanism by which Mettl3 regulates pausing, we mapped m^6^A modifications by methylated RNA immunoprecipitation followed by sequencing (MeRIP-seq), using a cell number-normalization approach (fig. S3A-E, and see Methods). We identified 15,046 m^6^A peaks within 7,095 genes, which represents 48% of all genes expressed in control or paused ESCs (Table S3). Principal Component Analysis (PCA) revealed that paused ESCs are in a distinct state with regards to the m^6^A RNA profile (Fig. 3A). Consistent with the quantitative mass spectrometry analysis (Fig. 1A), MeRIP-seq showed a global increase in m^6^A in paused ESCs, with 1,562 peaks significantly hypermethylated versus only 249 regions being hypomethylated relative to control ESCs (Fig. 3B-C, fig. S3F-G).

**Fig. 3.**
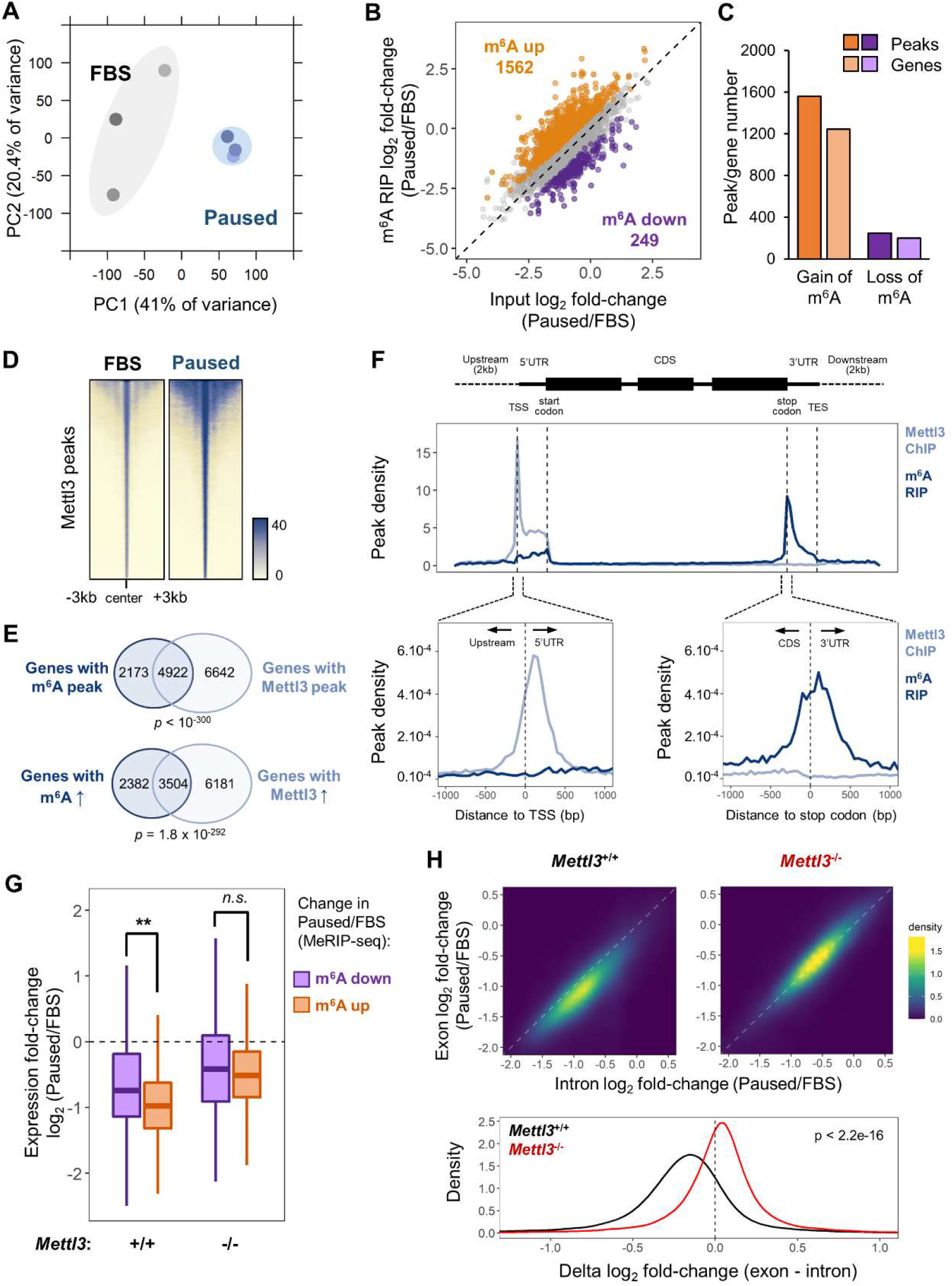
Mettl3-mediated m_6_A promotes RNA instability during pausing. (**A**) PCA plot showing that paused ESCs have distinct m^6^A profiles (n=3). (**B**,**C**) MeRIP-seq shows increased m^6^A in paused ESCs (n=3, gain/loss: adjusted *P* < 0.05 and absolute fold-change >1.5). (**D**) Heatmaps of Mettl3 ChIP-seq signal in FBS-grown and paused ESCs, showing increased Mettl3 binding in paused ESCs (n=2). (**E**) Overlap between targets of m^6^A and Mettl3, identifying all target genes or genes with increased levels of m^6^A and Mettl3 (log_2_FC > 0) in paused ESCs. *P*-value by hypergeometric test. (**F**) Metagene profiles of m^6^A and Mettl3 peaks. (**G**) RNAs with increased m^6^A in pausing, as defined in (A), are significantly more downregulated than RNAs with decreased m^6^A in paused *Mettl3*^+/+^ (but not *Mettl3*^-/-^) ESCs. Student ’s *t*-tests \*\**P* < 0.01. (**H**) Differences in expression (log_2_FC paused/FBS) between exonic and intronic RNA-seq data indicate a global decrease in RNA stability in *Mettl3*^+/+^ (but not *Mettl3*^-/-^) ESCs upon pausing. *P*-value by paired Student ’s *t*-test.

To understand the gain in m^6^A in paused ESCs, we investigated the levels of Mettl3 protein. Interestingly, despite no change in whole cell levels, Mettl3 is increased in the chromatin compartment upon transition to the paused state (fig. S4A-B). Mettl3 has previously been shown to bind to chromatin in ESCs and cancer cells, where it deposits m^6^A co-transcriptionally(*15, 16*). We therefore mapped the genome-wide distribution of Mettl3 by chromatin immunoprecipitation-sequencing (ChIP-seq) (fig. S3C-D). As anticipated, we observed higher levels of Mettl3 occupancy in paused ESCs relative to control ESCs (Fig. 3D). Although Mettl3 binds extensively throughout the genome, it is more abundant over expressed genes, particularly if their mRNAs are also m^6^A methylated (fig. S4E). The majority of m^6^A methylated RNAs (4,922/7,095, 69.4%) are transcribed from genes occupied by Mettl3 in ESCs, and transcripts gaining m^6^A during pausing largely arise from genes with elevated Mettl3 binding in paused ESCs (3,505/5,886, 59.6%, Fig. 3E). In agreement with recent reports(*15, 17*), Mettl3 localizes mainly to the transcriptional start site (TSS), while m^6^A is enriched near the stop codon and 3 ’ untranslated region (UTR) of coding genes in ESCs (Fig. 3F). Overall, these results indicate that an increased chromatin recruitment of Mettl3 underlies the gains of m^6^A RNA methylation during paused pluripotency, although we cannot exclude that Mettl3 may also exert other functions at chromatin.

To dissect the function of m^6^A RNA methylation during pausing, we asked how changes in m^6^A impact mRNA levels during ESC pausing. In the context of global hypotranscription in wildtype ESCs upon pausing ((*5*) and Fig. 2F), RNAs with increased m^6^A are significantly more downregulated than RNAs with decreased m^6^A (Fig. 3G). However, this association is lost in *Mettl3*^-/-^ cells, when analyzing the same genes (Fig. 3G). These data suggest that Mettl3-mediated m^6^A RNA methylation may contribute to transcript destabilization during pausing. In order to further explore this possibility, we re-analyzed the RNA-seq data from control and paused *Mettl3*^+/+^ and *Mettl3*^-/-^ ES cells to assess post-transcriptional regulation (Fig. 2F and fig. S2). Exonic reads reflect steady-state mature mRNAs, whereas intronic reads mostly represent pre-mRNAs, and therefore comparing the difference between these has been shown to effectively quantify post-transcriptional regulation of gene expression(*18*) (fig. S5A-C). This analysis pointed to a global destabilization of the transcriptome when *Mettl3*^+/+^ ESCs are induced to the paused state (Fig. 3H). In contrast, this global destabilization effect of pausing is entirely lost in *Mettl3*^-/-^ ESCs (Fig. 3H). Taken together, these results indicate that Mettl3-dependent m^6^A methylation is responsible for a global destabilization of the transcriptome in paused ESCs.

Our findings to this point indicate that the transcriptionally dormant state of paused cells is imparted by a combination of reduced nascent transcription and increased transcript destabilization, effects that are muted in *Mettl3*^-/-^ paused ESCs (Fig. 2C, fig. S5D-E). We hypothesized that m^6^A may contribute to destabilizing mRNAs encoding putative “anti-pausing” factors. To identify such factors, we mined the RNA-seq and MeRIP-seq data for genes that i) gain m^6^A in paused ESCs; ii) are downregulated upon pausing but to a lesser extent in *Mettl3*^-/-^ ESCs; iii) are expressed at least 2× higher in paused *Mettl3*^-/-^ ESCs relative to control paused ESCs; and iv) are destabilized in paused ESCs in a Mettl3-dependent manner. This analysis identified 953 candidate anti-pausing factors (fig. S5F, see Methods). We then took advantage of published data from early mouse embryos(*4*) to rank the candidates by their correlation with an expression signature of the m^6^A machinery. We reasoned that, if these candidate genes are regulated by m^6^A in vivo, their expression should be anti-correlated with expression of the methyltransferase complex and correlated with expression of the m^6^A demethylation machinery (Fig. 4A, Table S5, see Methods for details). Remarkably, the top-ranked candidate from this analysis is *Mycn*, which codes for the N-myc proto-oncogene (N-Myc) and is highly expressed in both ESCs and embryos (fig. S6A). The Myc-regulated set of genes is a major module of the ESC pluripotency network and it is downregulated in diapause(*4, 13*). Importantly, Myc family members can act as global transcriptional amplifiers in the context of development and cancer(*19, 20*). We therefore investigated in-depth the regulation of *Mycn* by m^6^A RNA methylation and its potential impact in paused ESCs.

**Fig. 4.**
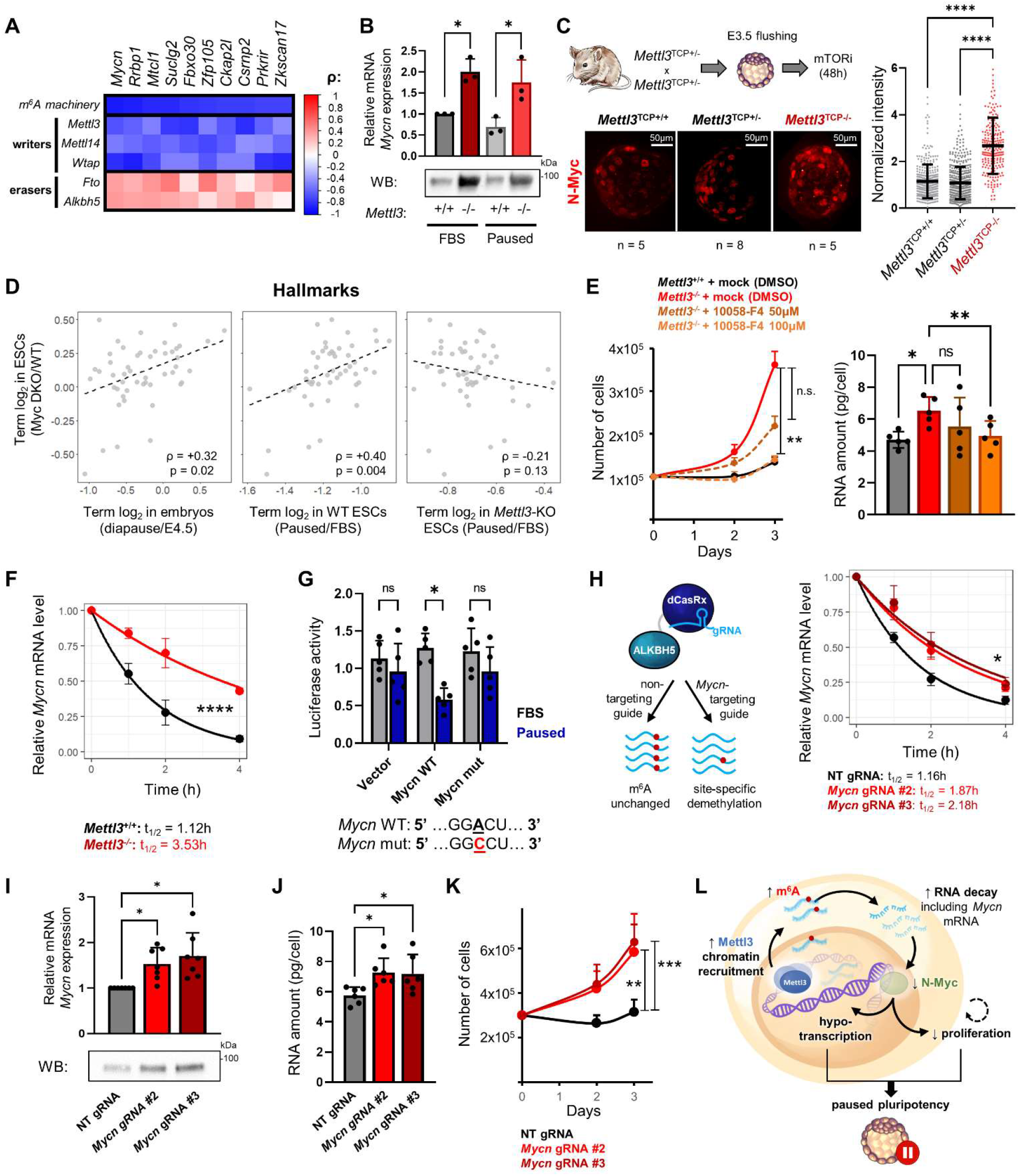
Mettl3 regulates pausing via m_6_A-mediated destabilization of *Mycn* mRNA. (**A**) Top putative anti-pausing factors ranked by Spearman correlation (ρ) with the m^6^A machinery. (**B**) Increased N-Myc in *Mettl3*^-/-^ ESCs by RNA-seq (n=3) and western blot (n=4). (**C**) Immunofluorescence and nuclear signal quantification of N-Myc in ex vivo paused blastocysts. Median log_2_ fold-changes for “hallmark” gene sets in *Myc/Mycn* DKO ESCs versus diapaused embryos or paused ESCs, with spearman correlation (ρ). (**E**) Blocking Myc signaling partially restored the decreases in proliferation (n=4) and total RNA (n=5) in paused *Mettl3*^-/-^ ESCs. (**F**) Increased *Mycn* mRNA stability in paused *Mettl3*^-/-^ ESCs (n=3). t_1/2_: half-life. (**G**) *Mycn* m^6^A site reduces transcript stability in paused ESCs (n=5). (**H**-**K**) Site-specific demethylation of *Mycn* leads to increased *Mycn* mRNA stability (H, n=3). increased expression (I, n=7 by RT-qPCR and n=3 by western blot), increased total RNA (J, n=6) and higher proliferation (K, n=5). (**L**) Model for the regulation of paused pluripotency by m^6^A RNA methylation. All data are mean ± SD, except times series which are mean ± SEM. *P*-values by two-tailed paired Student ’s *t*-tests (B,C,G), linear regression test (E left,F,H,K), one-way ANOVA with Dunnett ’s multiple comparison tests (E right,I,J). \**P* < 0.05, \*\**P* < 0.01, \*\*\**P* < 0.001, \*\*\*\**P* < 0.0001.

We found that N-Myc expression is elevated at the RNA and protein level in paused *Mettl3*^-/-^ ESCs and blastocysts (Fig. 4B-C, fig. S6B-D), further supporting its status as an m^6^A target. Additionally, ESCs depleted for both c-Myc and N-Myc (Myc DKO) partially recapitulate gene expression changes in diapaused embryos and paused ESCs(*5, 13*), but this relationship is entirely abolished in paused *Mettl3*^-/-^ ESCs (Fig. 4D). In agreement with this result, the downregulation of Myc target genes that occurs upon pausing is partially suppressed in *Mettl3*^-/-^ ESCs (Fig. 2H). We thus wondered whether elevated Myc signaling contributes to the defective pausing observed in *Mettl3*^-/-^ ESCs. Accordingly, treatment with the Myc inhibitor 10058-F4 restored the decreases in proliferation and total RNA content in paused *Mettl3*^-/-^ ESCs to levels equivalent to paused *Mettl3*^+/+^ cells (Fig. 4E, fig. S6E). The downregulation of Myc target genes in pausing is imparted via reduced nascent transcription, rather than by being directly targeted by m^6^A themselves (fig. S6F-G), consistent with the established role of Myc as a transcriptional activator(*19, 20*). Taken together, these results indicate that N-Myc levels are downregulated in an m^6^A-dependent manner in paused ESCs, and that the subsequent decrease in Myc signaling results in reduced transcriptional output and proliferation.

We sought to further probe the direct regulation of *Mycn* mRNA by methylation. The m^6^A mark can affect mRNA metabolism through binding of reader proteins, including the YTH domain family and HNRNP proteins(*21*). In particular, the reader Ythdf2 can mediate destabilization of m^6^A methylated mRNAs, including in ESCs. In agreement with this, *Mycn* mRNA is bound by both Mettl3 and Ythdf2 in paused ESCs, and this binding is abolished by knockout of *Mettl3* (fig. S7A). Moreover, the half-life of *Mycn* mRNA is significantly increased in *Mettl3*^-/-^ ESCs (Fig. 4F), specific to *Mycn* among the Myc family members (fig. S7B-C). These results corroborate that m^6^A methylation regulates *Mycn* mRNA stability in paused ESCs. After identifying a hypermethylated m^6^A site near the stop codon of the *Mycn* mRNA (fig. S8A-B), we found that this site confers transcript destabilization in paused ESCs in a luciferase reporter assay, in a manner dependent on the integrity of the m^6^A site as an A to C mutation nullifies this effect (Fig. 4G).

Finally, we explored how RNA methylation impacts *Mycn* transcript stability. We performed site-specific RNA demethylation using dCasRx-conjugated ALKBH5(*22*) (fig. S8C-G). We found that targeted demethylation of the 3 ’ end of *Mycn* mRNA in *Mettl3*^+/+^ paused ESCs stabilizes the transcript, leading to higher steady-state levels of N-Myc mRNA and protein (Fig. 4H-I, fig. S8H). Remarkably, this targeted loss of m^6^A at the *Mycn* mRNA is sufficient to increase levels of total RNA and proliferation in paused ESCs (Fig. 4J-K). Thus, m^6^A demethylation of *Mycn* mRNA in otherwise wildtype ESCs recapitulates the suppression of pausing observed in *Mettl3*^-/-^ ESCs.

In summary, we show that Mettl3-dependent m^6^A RNA methylation is required for developmental pausing by maintaining transcriptional dormancy (Fig. 4L). Our findings shed light on the molecular mechanisms that underlie mammalian developmental pausing and reveal Mettl3 as a key integrator between transcriptomic and epitranscriptomic levels of gene regulation. m^6^A RNA methylation was recently reported to modulate the transcriptional state of ESCs by destabilizing chromosome-associated RNAs and transposon-derived RNAs(*23–25*) and by promoting the recruitment of heterochromatin regulators(*17*). Our results support the notion that Mettl3 acts at chromatin and methylates RNA co-transcriptionally(*15, 16*). Surprisingly, even though Mettl3 methylates thousands of transcripts, we found that one target, *Mycn* mRNA, is key for its function in maintaining the suppressed transcriptional state of paused cells. Future studies may uncover additional functions of other m^6^A-regulated putative anti-pausing factors identified here.

We anticipate that the regulatory relationship between m^6^A RNA methylation and cellular dormancy will have implications extending well beyond embryonic diapause. Modulations of mTOR signaling have been implicated in the control of stem cell dormancy in various embryonic and adult tissues(*26*). Moreover, we and others have shown that cancer cells can enter a dormant state molecularly and functionally similar to diapause to survive chemotherapy(*27, 28*). The insights gained here provide exciting new opportunities to explore the biology of m^6^A RNA methylation in the fields of developmental biology, reproductive health, regenerative medicine, and cancer.

## Acknowledgments

We thank members of the Santos Lab, D. Schramek, A. Bulut-Karslioglu, T. Macrae, J. Jeschke and F. Fuks for feedback on the manuscript. We are grateful to the Hanna lab for providing cells, to the Toronto Center for Phenogenomics for mouse colony maintenance and support, members of the UCSF Center for Advanced Technology and the LTRI Sequencing Core for next-generation sequencing, the LTRI Flow Cytometry Facilities, A. Bulut-Karslioglu and S. Biechele for advice on diapause, M. Percharde, T. Macrae and S. McClymont for bioinformatics guidance.

## Funding

Belgian American Educational Foundation Inc (EC).

US National Institutes of Health GM58843 (PAL)

US National Institutes of Health R01GM113014 (MRS)

Canada 150 Research Chair in Developmental Epigenetics (MRS)

Great Gulf Homes Charitable Foundation (MRS)

Canadian Institutes of Health Research Project Grant 165935 (MRS)

Canadian Institutes of Health Research Project Grant 178094 (MRS)

## Data and materials availability

Sequencing data have been deposited on the NCBI Gene Expression Omnibus repository (GEO, http://ncbi.nlm.nih.gov/geo) and will be accessible upon publication. Published RNA-seq data used in this study are available under the accession numbers E-MTAB-2958 (early mouse embryos) and E-MTAB-3386 (*Myc/Mycn* DKO ESCs). All data are available in the main text or the supplementary materials.

## Supplementary Materials

Materials and Methods

Figs. S1 to S10

Tables S1 to S6

## Materials and Methods

### Mouse embryonic stem cell culture

E14 ESCs were grown on gelatin-coated plates in standard serum/LIF medium: DMEM GlutaMAX with Na Pyruvate, 15% FBS (Atlanta Biologicals), 0.1 mM Nonessential amino acids, 50U/ml Penicillin/Streptomycin, 0.1mM EmbryoMax 2-Mercaptoethanol, and 1000 U/ml ESGRO LIF. For FBS/2i culture, the medium was supplemented with 1μM PD0325901 and 3μM CHIR99021. Pausing was induced by adding 200nM INK128 to the medium, as described(*5*). Unless otherwise stated, ESCs were paused for at least 5 days before use, and for exactly 2 weeks for all sequencing experiments.

### *Mettl3*^*-/-*^ ESC models

*Mettl3*^-/-^ and control *Mettl3*^+/+^ ESCs were kindly provided by J. Hanna(*8*) and used for most experiments, except where indicated. An independent ESC line mutant for *Mettl3* (*Mettl3*^-/-#2^) cells was generated via CRISPR/Cas9 for validation of key results. Cloning was performed by annealing oligos targeting *Mettl3* into pSpCas9(BB)-2A-GFP (PX458), a gift from Feng Zhang (Addgene plasmid #48138; RRID:Addgene_48138)(*29*). E14 were transfected with lipofectamine 2000, isolated by FACS, clonally expanded and validated for Mettl3 loss.

### Site-specific m^6^A demethylation

Lentivirus was produced by transfecting HEK293T cells with pMSCV-dCasRx-ALKBH5-PURO and viral packaging/envelope vectors pMD2.G and psPax2, gifts from Qi Xie and Didier Trono (Addgene plasmid #175582, #12259 and #12260; RRID:Addgene_175582; RRID:Addgene_12259, RRID:Addgene_12260)(*22*). E14 cells were infected with pMSCV-dCasRx-ALKBH5-PURO lentivirus, and after 48h were selected with 2µg/mL puromycin. Clonal lines were expanded and selected for expression of dCasRx-ALKBH5 by western blot. Guide RNAs were cloned using lenti-sgRNA-BSD, a gift from Qi Xie (Addgene plasmid #175583; RRID:Addgene_175583)(*22*). Each guide plasmid (1μg) was transfected with Lipofectamine 2000 into dCasRx-ALKBH5-expressing cells in a 6-well plate. Cells were selected with 8μg/mL blasticidin for 3 days before use. See Table S6 for primer sequences.

### Mouse models

The mouse C57BL/6NCrl-Mettl3^em1(IMPC)Tcp^ allele was generated as part of the Knockout Mouse Phenotyping Program (KOMP2) project at The Centre for Phenogenomics (TCP) by Cas9-mediated deletion of a 142 bp region (Chr14:52299764 to 52299905 in ENSMUSE00001224053, GRCm38), causing a frameshift and early truncation (I171Mfs*4). Mice were purchased from the Canadian Mouse Mutant Repository through TCP. Heterozygotes mice (referred to as “*Mettl3*^TCP+/-^”) were maintained on a hybrid C57BL/6×CD-1 background in order to increase litter size and ex vivo pausing efficiency. Genotyping of mice was performed by Transnetyx. All mice were housed at TCP in Toronto. All procedures involving animals were performed according to the Animals for Research Act of Ontario and the Guidelines of the Canadian Council on Animal Care. All procedures conducted on animals were approved by The Animal Care Committee at TCP (Protocol 22-0331). Sample size was not predetermined.

For all embryo experiments, 6 to 12-week-old *Mettl3*^*TCP* +/-^ females were mated with 6-week-to 8-month-old *Mettl3*^*TCP* +/-^ males. Collection and ex vivo culture of blastocysts was performed as previously described(*5*), with pausing induced by flushing blastocysts at E3.5 and culturing them at 5% O2, 5% CO2 at 37 °C in KSOM^AA^, with addition of 200nM RapaLink-1 on the day after flushing. Blastocysts with collapsed blastocoel were considered non-viable and collected for genotyping every day. Hormonal diapause was induced as previously described(*5*), and blastocysts were collected at EDG8.5 and genotyped. For genotyping of embryos, DNA was extracted from individual blastocysts using the Red Extract-N-Amp kit (Sigma), in 36μl final volume. *Mettl3* status was assessed by PCR using 1μl DNA extract in 15μl total volume reaction with Phire Green Hot Start II PCR Master Mix (Thermo Fisher). Cycling conditions: 98°C for 30 s; 35 cycles of 98°C for 5s, 57°C for 5s, 72°C for 5s; 72°C for 1 min. See Table S6 for primer sequences.

### Cell number-normalized (CNN) RNA analyses

RNA was extracted from equal number of ESCs, typically ∼2×10^5^ cells, with RNeasy Micro Kit with on-column DNase I digestion (QIAGEN). RNA concentration was measured by Qubit™ RNA High Sensitivity to calculate the amount of total RNA per cell. cDNAs were generated using equal volumes of extracted RNAs with the SuperScript IV VILO Master Mix (Thermo Fisher) and qPCR data were acquired on a QuantStudio 5 (Thermo Fisher Scientific). Unless otherwise stated, gene expression was normalized according to cell number(*19*). See Table S6 for primer sequences.

### CNN RNA-sequencing and data analysis

RNA extracted from equal number of ESCs was spiked with synthetic RNAs from the External RNAs Control Consortium (ERCC) Spike-in Mix1 (Thermo Fisher), by adding 2µl of 1:100 ERCC dilution to 10µl of RNA (equivalent to ∼1-2µg). Library preparation was done using the NEBNext Ultra II Directional Library Prep Kit for Illumina with the mRNA Magnetic Isolation Module from 1µg RNA, per manufacturer ’s instructions (NEB). Library quality was assessed by Fragment Analyzer NGS (Agilent). Sequencing was performed on a NextSeq500 (Illumina) with 75bp single-end reads at the Lunenfeld-Tanenbaum Research Institute Sequencing Facility.

Libraries were trimmed of adaptors and quality-checked using Trim Galore! v0.4.0, and then aligned to the mm10 transcriptome with ERCC sequences using TopHat2 v2.0.13. Gene counts were obtained from the featureCounts function of subread (v1.5.0) with options: -t exon -g gene_id. Raw counts were imported into R, and normalized to ERCCs with edgeR (v3.32.1) as previously described(*19, 30*). Data were further analyzed using tidyverse v1.3.0 and plotted using ggplot2 v3.3.5. The threshold for significant differential expression was adjusted *P* < 0.05 and absolute fold-change > 1.5. Normalized counts were mean-centered per batch and log-transformed for PCA and heatmaps. GSEA was performed using the fGSEA package v1.16.0, pre-ranking genes by *t*-values from the differential analysis (paused/FBS). Gene set collections were downloaded from the Molecular Signatures Database v7.5.1 (http://www.gsea-msigdb.org/gsea/msigdb/index.jsp). For intron analysis, RefSeq-annotated intronic regions were shortened by 50bp on each side to limit overlap with exonic regions, then counts were obtained with featureCounts and analyzed in R as for exons.

### Global m6A quantification

For quantification of m^6^A in poly(A) RNA, nucleoside digestion was performed as previously described(*31*). Separation was accomplished by reversed phase chromatography with an Acquity UPLC HSS T3 (Waters) on a Vanquish™ Flex Quaternary UHPLC system (Thermo Fisher Scientific). Mass spectrometry was performed on a Quantiva™ triple quadrupole mass spectrometer interfaced with an H-ESI electrospray source (Thermo Fisher Scientific). Data were analyzed with Tracefinder 4.1 (Thermo Fisher Scientific) and Qual browser of Xcalibur 3.0. The mass transitions (precursor → product) for m^6^A were 282 → 94, 282 → 123 and 282 → 150. Changes in m^6^A were also measured by dot blot from 50ng of poly(A) RNA. Blotting was performed as previously described for 5-hydromethylcytosine(*32, 33*), except that Diagenode C15410208 (1:400) was used as primary antibody.

### m^6^A MeRIP-seq

ESCs were spiked with 2% of human cells (HeLa, CVCL_0030), then poly(A) RNA was extracted using the Magnetic mRNA Isolation Kit (NEB). Immunoprecipitation of methylated RNA (MeRIP) was done using the EpiMark® N6-Methyladenosine Enrichment Kit with 4µg spiked poly(A) RNA. Specificity of the IP was tested by spiking samples with exogenous RNA controls (m6A modified and unmodified), per manufacturer ’s instructions. The cell number normalization approach with human cells was validated beforehand by spiking ESCs with 1, 2 or 4% human cells to simulate global changes in methylation (see fig. S3b). m^6^A enrichment was measured by RT-qPCR in 3 mouse mRNAs (*Neurod1, Nr5a2, Sox1*) known to be methylated in ESCs(*34*) and normalized to the average levels of 5 highly-expressed and methylated human mRNAs (*HSBP1, PCNX3, GBA2, ITMB2, PCBP1*)(*35*). MeRIP libraries were constructed from 0.5-1ng of input or IP RNA and prepared using the SMARTR-seq RNA library prep v2 kit (TakaraBio), per manufacturer ’s recommendations. Library quality was assessed with the High Sensitivity DNA Assay on an Agilent 2100 Bioanalyzer (Agilent Technologies), and samples were sequenced on a HiSeq 4000 using single-end 50 bp reads at the UCSF Center for Advanced Technology.

Pre-processing of sequencing data was performed similarly to RNA-seq, but without ERCCs and with reads unmapped to mm10 being aligned to hg19. For input samples, gene expression was normalized as for CNN RNA-seq, except that the ratio of hg19/mm10 reads was used for normalization instead of ERCCs. For m^6^A RIP samples, peaks were called with MACS2 (using IP samples and their input counterpart as controls and q<0.01). Peaks were annotated by intersecting center positions with RefSeq annotations. Peak analysis was performed using DiffBind v3.0.15, with the options minOverlap=2, score=DBA_SCORE_READS. MeRIP peaks were then first normalized using the ratio of hg19/mm10 reads in each sample for normalization, then adjusted by dividing values by the ratio Input_sample_/Input_average_ of the corresponding gene to consider expression changes. In the following differential analysis with edgeR, these normalized m^6^A levels were protected from further re-scaling by fixing the library size for all samples as lib.size = rep(10^6, 6) in the voom function. The threshold for significant differential methylation was adjusted *P* < 0.05 and absolute fold-change >1.5. For motif analysis, all peaks were limited to 100bp surrounding the center and submitted to DREME of the Meme-suite (http://meme-suite.org). Bigwig files were generated by Deeptools v3.3.0 and visualized in Integrated Genome Viewer (IGV v2.9.4), with the vertical scale being adjusted to consider expression changes individually for each gene (similar to the m^6^A value adjustment performed before the differential analysis).

### Site-specific quantification of m^6^A

We identified a putative m^6^A site within the *Mycn* MeRIP peak using the m^6^A-Atlas database (http://www.xjtlu.edu.cn/biologicalsciences/atlas)(*36*). We then measured m^6^A by RT-qPCR, following a method previously described(*37, 38*), which takes advantage of the diminished capacity of Bst enzymes to retrotranscribe m^6^A residues compared to MRT control enzyme, and RT primers targeting just before or after the site (primer + or −). Each cDNA was generated with ∼100ng of total RNA, 100nM primer (+ or −), 50μM dNTPs and 0.1U of Bst3.0 (NEB) or 0.8U of MRT (ThermoScientific). The cycling conditions were 50°C for 15min, 85°C for 3min, then 4°C. RT-qPCR data were then acquired on a QuantStudio 5 (Thermo Fisher Scientific) and normalized as [2^-(Ct_Bst−_ - Ct_MRT−_) -2^-(Ct_Bst+_ - Ct_MRT+_)] / 2^-(Ct_Bst−_ - Ct_MRT−_). Negative values were considered below the detection threshold and rounded to 0.

### Mettl3 ChIP-seq

ESCs were spiked with 2% of human cells (HeLa), then cross-linked in 1% formaldehyde/PBS for 10min at room temperature. After quenching with 125mM glycine for 5min at room temperature, followed by 15min at 4°C, cells were washed in cold PBS and stored at −80 °C. Cells were diluted at 5 million cells per 100μl in shearing buffer (1% SDS, 10mM EDTA, 50mM Tris-HCl pH8.0, 5mM NaF, Halt™ Protease Inhibitor Cocktail (Thermo Fisher), 1mM PMSF), rotating at 4°C for 30min, then 100μl of lysate was passed into a microTUBE AFA Fiber Snap-Cap (Covaris). Chromatin was sheared to 200–500 bp fragments on a Covaris E220 with settings PIP 175, Duty 10%, CPB 200, for 7min. Immunoprecipitation was performed overnight using 200μl of each lysate (∼chromatin from 10 million cells) and 5µg of antibody (Proteintech 15073-1-AP), following the iDeal ChIP-seq kit for Transcription Factors (Diagenode) protocol. Elution, de-crosslinking, and DNA purification were performed per manufacturer ’s instructions. Libraries were constructed from ∼2ng DNA and prepared using the NEBNext Ultra II DNA Library Prep Kit for Illumina (NEB). Library quality was assessed by Fragment Analyzer NGS. Samples were sequenced on a NextSeq500 (Illumina) with 75bp single-end reads at the Lunenfeld-Tanenbaum Research Institute Sequencing Facility.

Reads that passed quality control were trimmed of adaptors using Trim Galore! v0.4.0 and aligned to mm10 using bowtie2 v2.2.5131. Unmapped reads were then mapped to hg19. SAM files were converted to BAM files, sorted, and indexed with samtools v1.9. Bam files were deduplicated using MarkDuplicates (picard v2.18.14). Peaks were called with MACS2 (using IP samples and their input as controls) with the options --gsize 3.0e9 -q 0.05 –nomodel --broad and annotated by intersecting center positions with RefSeq annotations. The most upstream and downstream annotated TSS and TES, respectively, were considered for each gene. Peak analysis was performed using DiffBind v3.0.15 with score=DBA_SCORE_TMM_READS_FULL_CPM. Normalization was done with edgeR, using the ratio of mm10/hg19 reads (relative to respective input sample). For TSS analysis, a 1kb window surrounding the TSS of every RefSeq gene was used. Bigwig files were generated with Deeptools v3.3.0 using –scaleFactor for normalisation and visualized in Integrated Genome Viewer (IGV v2.9.4).

### Screening for m^6^A targets

First, we selected all genes that gain m^6^A in pausing (MeRIP-seq: log_2_FC > 0). Second, we looked for genes whose downregulation is suppressed in *Mettl3*^-/-^ ESCs (RNA-seq: logFC_*Mettl3*+/+_ < 0 & log_2_FC_*Mettl3*+/+_ < log_2_FC_*Mettl3*-/-_). Third, we looked for genes whose expression is at least 2× higher in paused *Mettl3*^-/-^ ESCs (RNA-seq: CPM_*Mettl3*-/-_ > 2× CPM_*Mettl3*+/+_). Finally, we selected genes whose RNA is more stable in paused *Mettl3*^-/-^ ESCs (RNA-seq: log_2_FC_exon_ > 1.5 × log_2_FC_intron_). We then used published RNA-seq data from early mouse embryos(*3*), with expression data first transformed into z-scores. To establish a quantitative signature of the m^6^A machinery, we averaged the z-scores of the writers (*Mettl3, Mettl14, Wtap*) and the z-scores of the erasers (*Fto, Alkbh5*) multiplied by −1. We then ranked all targets by their Spearman correlation coefficients with the m^6^A machinery signature, focusing on genes with negative correlations, reflecting a potential destabilization of mRNAs in presence of the m^6^A machinery in embryos.

### RNA immunoprecipitation

For RIP experiments, 2µg anti-Mettl3, Ythdf2 or control IgG antibodies were pre-bound to 20µL Protein A Dynabeads (Thermo Fisher), rotating for 3h at 4°C. Beads were next collected on a DynaMag (Thermo Fisher) and resuspended in RIP buffer (150mM KCl, 25mM Tris pH7.4, 5mM EDTA, 0.5mM DTT, 0.5% NP40, protease and RNase inhibitors) containing 500ng/mL tRNA (Thermo Fisher) and 1mg/mL BSA to block for 30 min. During incubation time, ESCs were collected and lysed in RIP buffer on ice for 20min. Supernatants (500µl, equivalent to 10 million cells) were pre-cleared with 20µL Protein A Dynabeads, rotating for 30min at 4°C. Cleared lysates were incubated together with antibody-bound blocked beads overnight at 4°C. The next day, lysates were washed 5 times in RIP buffer, and RNA was extracted using Direct-zol RNA Kits (Zymo Research), before analyzing by RT-qPCR.

### Western blot analysis

Whole-cell and chromatin extract we prepared according to methods described previously(*39, 40*). Denatured samples were separated on 4-15% Mini-Protean TGX SDS-PAGE gels (Bio-Rad) and transferred to nitrocellulose membranes by wet transfer using Trans-Blot® Turbo™ Mini Nitrocellulose Transfer Packs (Bio-Rad). Membranes were blocked in 5% milk/TBS-T and incubated with indicated primary antibodies overnight at 4 °C. HRP-conjugated anti-mouse/rabbit secondary antibodies, were incubated for 1h at room temperature. Proteins were detected by ECL (Pierce) or Clarity Max (Bio-Rad). Quantification of bands was performed using ImageJ. See Table S7 for antibodies and dilutions.

### mRNA stability assay

Cells were treated with 5μg/ml actinomycin D (Sigma-Aldrich) for 0, 1, 2 or 4h before cell collection. RNA level was measured by RT-qPCR and normalized to *Actb*, a highly stable transcript. Expression values (relative to 0h) were fitted to an exponential decay model using linear regression in R.

### Luciferase reporters of RNA stability

The region surrounding the stop codon of *Mycn* mRNA, including the identified m^6^A site (wildtype sequence or A-to-C mutation) was cloned into the pmirGLO Dual-Luciferase miRNA Target Expression Vector (Promega), per manufacturer ’s instructions. Vector (500ng) was transfected in ESCs with lipofectamine 2000. After 48h, cells were lysed and Firefly luciferase signal was measured with a luminometer and normalized to Renilla luciferase activity, using Dual-Glo® Luciferase Assay System.

### Nascent transcription assays in ESCs

To assess global transcriptional output, cells were treated with 1mM 5-ethynyl uridine (EU) for 45min, collected by trypsinization and prepared following the Click-iT™ RNA Alexa Fluor 488 Imaging Kit (Thermo Fisher) instructions. Data were collected by flow cytometry using the Beckman Coulter Gallios and analyzed using Kaluza. Fluorescence values were plotted as median fluorescence intensity (MFI) per sample, relative to FBS *Mettl3*^+/+^ cells. For nascent RNA capture, EU incubation was performed in ESCs as above. Cells were collected by trypsinization, counted, and 2×10^5^ were used to extract RNA. Biotinylated nascent RNA was captured according to protocols within the Click-iT Nascent RNA Capture Kit (Invitrogen) and used in RT-qPCR.

### Embryo immunofluorescence

Ex vivo paused embryos were labelled in their culture medium for 45min with 1mM EU for nascent transcription, then fixed in 4% paraformaldehyde for 15 min. Permeabilization was done with 0.5% Triton X-100 in PBS + 5% FBS for 15 min. After blocking in PBS, 2.5% BSA, 5% donkey serum for 1h, embryos were incubated overnight at 4°C with the primary antibodies (Mettl3 Abcam ab195352 1/200; N-Myc Cell Signaling Technology D1V2A 1/200). EU fluorescence coupling was performed following the manufacturer ’s instructions for Click-iT™ RNA Alexa Fluor 488 Imaging Kit. Embryos were incubated with fluorescence-conjugated secondary antibodies for 1h at room temperature. Embryos were stained with DAPI in fresh blocking, washed twice, and transferred into a drop of M2 media (∼5μl).

Images were captured using a Leica DMI 6000 Spinning Disc Confocal microscope, and embryos were genotyped as before. Quantification was performed with ImageJ, with 10 consecutive image planes (imaged at 10μm intervals) stacked by “average intensity” projection. This was repeated 4 times (totaling 40 image planes used per embryo). Then, individual nuclei were quantified using the ROI Manager, and the background (captured outside of the embryo) was subtracted to values within each projection. To avoid batch effects, values were also normalized to the average of *Mettl3*^TCP +/+^ embryos within each litter.

### Statistics and reproducibility

Statistical analyses were performed in GraphPad Prism v9.3.1 or R v4.0.3. Data are presented as mean ± SD or SEM, except where otherwise indicated. Box plots present center lines as medians, with box limits as upper and lower quartiles and whiskers as 1.5×IQR. Two-tailed Student ’s t-test and one- or two-way ANOVA with Dunnett ’s multiple comparison tests were used when normal distribution could be assumed. Time series were modeled by linear regression on log_2_-transformed y values, with *P*-values extracted from the interaction between time and the categorical variable of interest. GSEA was performed with fGSEA in R, with the adjusted *P*-value as indicated. Correlation was measured by *ρ* Spearman ’s rank correlation coefficient. All replicates for in vitro data are derived from independent experiments.

Sample size, number of replicates, errors bars and statistical tests were chosen based on experience and variability of in vitro studies and stated in each figure legend. Unless otherwise indicated, all experiments included technical replicates and were repeated at least three independent times. No statistical methods were used to predetermine sample sizes. All replicates for in vivo data are derived from at least 3 embryos per genotype and 2 separate litters. No statistical method was used to predetermine sample size. No data were excluded from the analyses. No randomization was required for design of in vivo experiments, as embryos were harvested, cultured and treated together and only later identified by genotype (Mettl3^+/+^, Mettl3^+/-^ or Mettl3^-/-^). No blinding was applied in this study, as all data produced derived from objective quantitative methods. To ensure consistent experimental conditions, all control and treatment samples were processed in parallel.

### Materials availability

Plasmids used in this study were generated from vectors commercially available from Addgene (#48138, #175582, #12259, #12260, #175583) and Promega (pmirGLO Dual-Luciferase miRNA Target Expression Vector). All sequences used for cloning are listed at Table S6.

### Data and code availability

Sequencing data have been deposited on the NCBI Gene Expression Omnibus repository (GEO, http://ncbi.nlm.nih.gov/geo) and will be accessible upon publication. Published RNA-seq data used in this study are available under the accession numbers E-MTAB-2958 (early mouse embryos) and E-MTAB-3386 (*Myc/Mycn* DKO ESCs). The authors declare that all other data supporting the findings of this study are available within the paper and its supplementary information files. Code supporting this study are available at a dedicated Github repository [https://github.com/EvelyneCollignon/Mettl3_pausing].

**Fig. S1.**
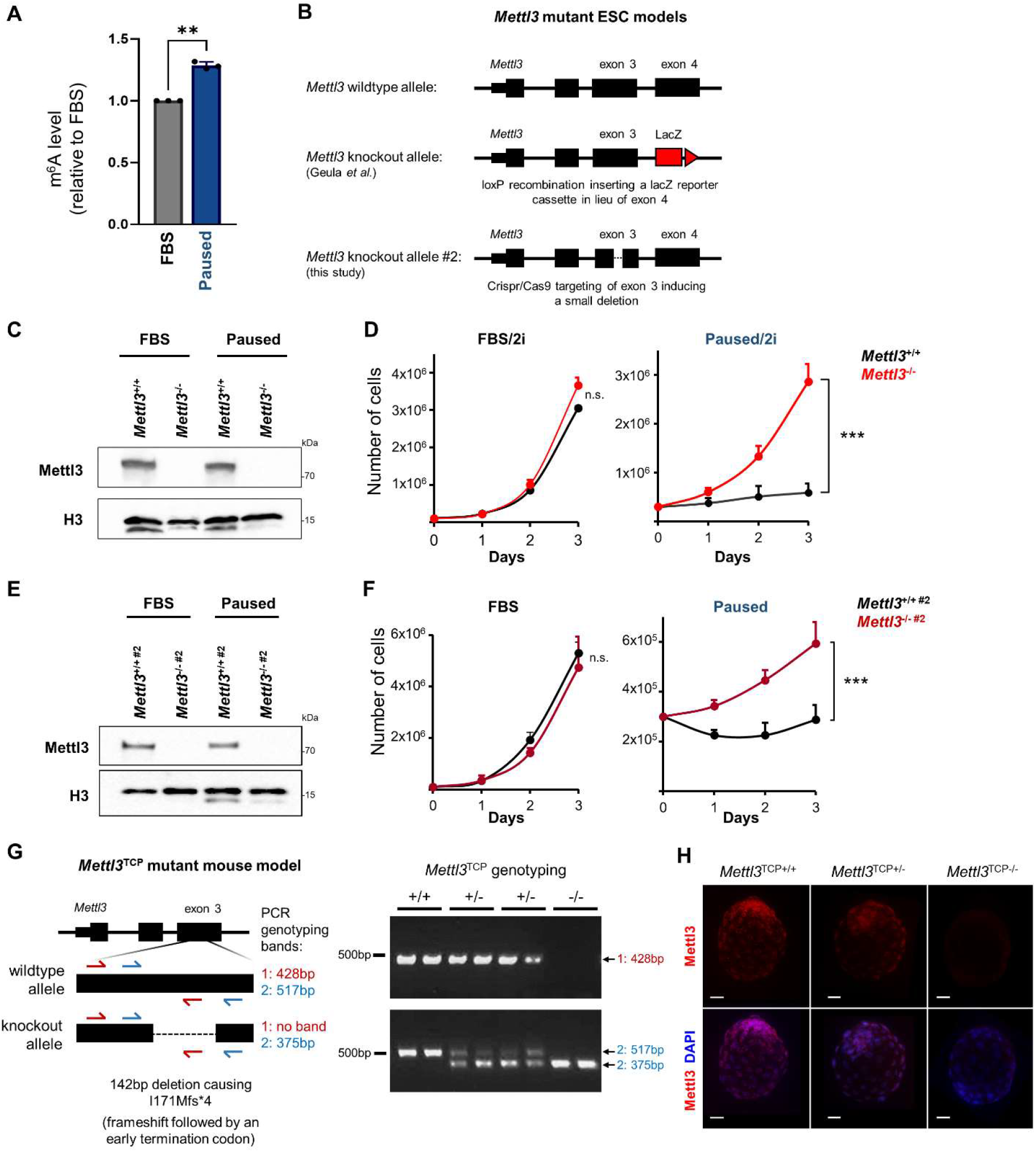
Dissection of paused pluripotency in *Mettl3*_-/-_ models. (**A**) m^6^A increase in paused ESCs was validated in an independent mass spectrometry experiment. Data are relative to FBS, as mean ± SD (n=3). Ratio paired Student ’s *t*-tests \**P* < 0.05. (**B**) Design of *Mettl3*^-/-^ ESCs models used in this study. (**C**) Validation of *Mettl3*^-/-^ ESCs, in FBS and pausing conditions, by western blot. (**D**) In FBS/2i culture, *Mettl3*^-/-^ ESCs also fail to suppress proliferation in paused conditions. Data are mean ± SEM (n=3). Linear regression test \*\*\**P* < 0.001. (**E**) Validation of *Mettl3*^-/-#2^ ESCs, in FBS and pausing conditions, by western blot. (**F**) *Mettl3*^-/-#2^ ESCs also fail to suppress proliferation in paused conditions. Data are mean ± SEM (n=3). (**G**) *Mettl3*-knockout mutant model in mice (*Mettl3*^TCP-/-^) with genotyping strategy and examples of PCR genotyping of embryos resulting from *Mettl3*^TCP+/-^ crossing. +/+: wildtype, +/-: heterozygous, -/-: knockout. (**H**) Validation of *Mettl3*^TCP-/-^ in embryos by immunofluorescence. Representative staining images are shown. Scale bars = 20 µm.

**Fig. S2.**
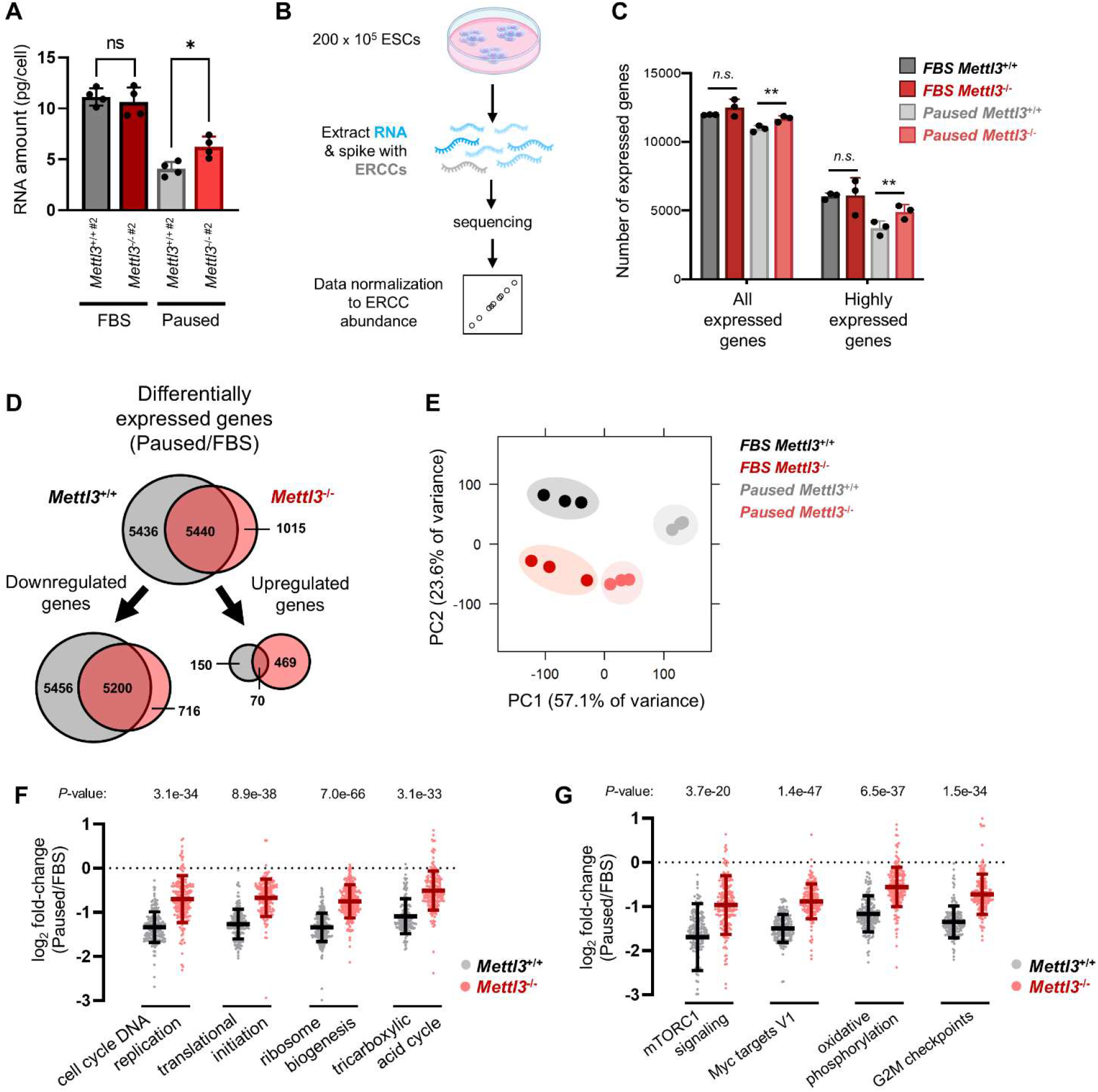
Paused *Mettl3*_-/-_ ESCs acquire a distinct gene expression profile. (**A**) Quantification of total RNA per cell in *Mettl3*^+/+ #2^ and *Mettl3*^-/-#2^ ESCs, in FBS and paused conditions. Data are mean ± SD (n=3). Student ’s *t*-tests \**P* < 0.05. (**B**) Strategy for RNA-seq with cell number-normalization using ERCC spike-in RNAs. (**C**) Quantification of the number of expressed genes in *Mettl3*^+/+^ and *Mettl3*^-/-^ ESCs, in FBS and paused conditions. Expressed genes are further defined as having high expression (log_2_ normalized reads > 5). Data are mean ± SD (n=3). Student ’s *t*-tests \*\**P* < 0.01. (**D**) Number of differentially expressed genes (fold-change > 1.5 and adjusted *P* < 0.05) upon pausing, in *Mettl3*^+/+^ and *Mettl3*^-/-^ ESCs. (**E**) PCA plot for all expressed genes across all samples, showing across PC1 that *Mettl3*^+/+^ ESCs acquire a more divergent expression profile upon pausing than *Mettl3*^-/-^ ESCs, relative to respective FBS condition. (**F**,**G**) Gene expression changes (log_2_ fold-changes) of gene sets selected from the “GO biological processes” (F) and “hallmarks” (G) collections, showing incomplete downregulation in paused *Mettl3*^-/-^ ESCs. Student ’s *t*-tests.

**Fig. S3.**
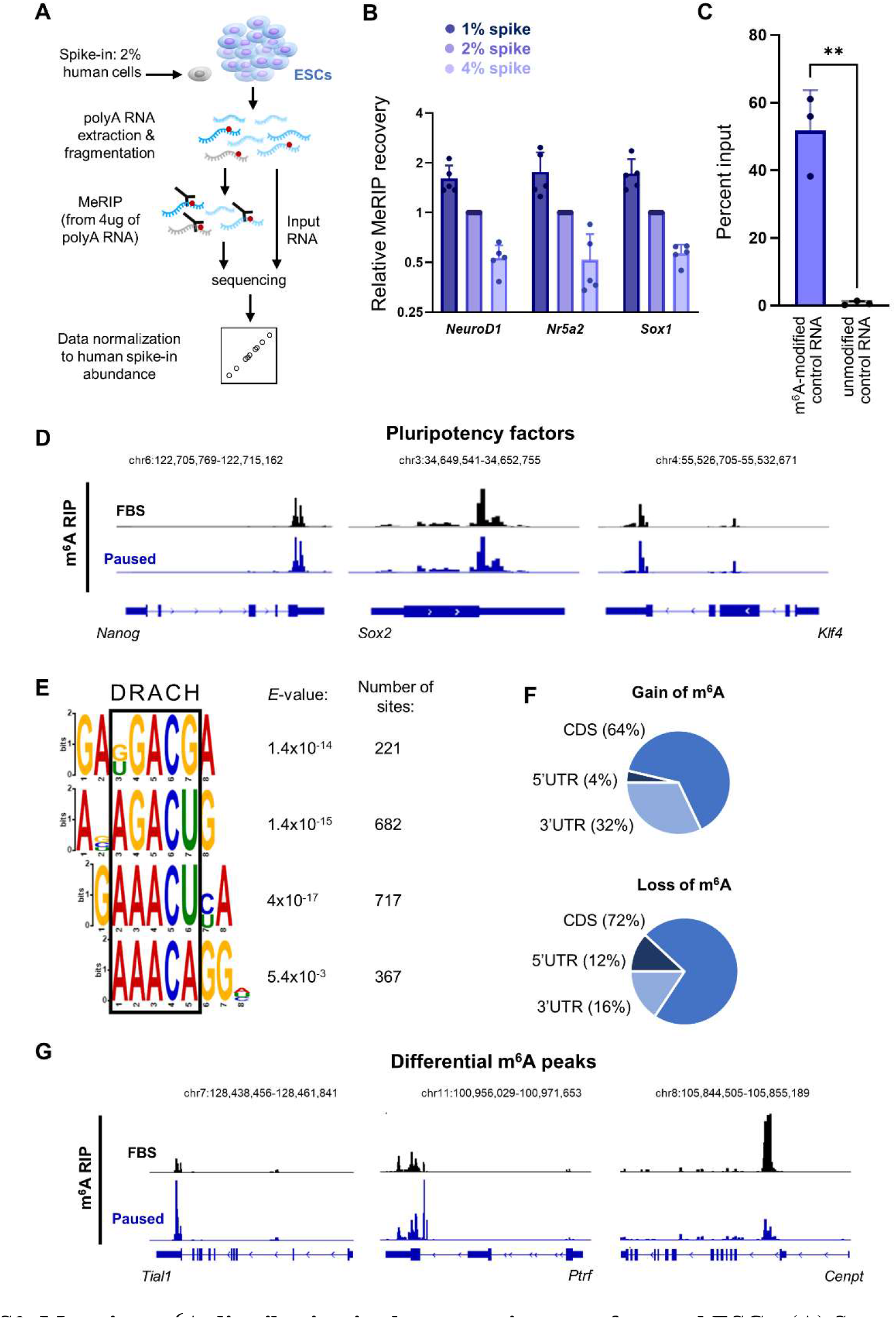
Mapping m_6_A distribution in the transcriptome of paused ESCs. (**A**) Strategy for MeRIP-seq in ESCs with cell number-normalization (CNN) using human cell spiking. (**B**) Validation of the CNN strategy for the MeRIP-seq. By mixing different ratios of human cells to ESCs (1, 2 or 4%), we simulated global changes in methylation. Spiking normalization allows capture of these differences, as shown here by MeRIP-qPCR for 3 methylated mRNAs (*NeuroD1, Nr5a2, Sox1*). Data are relative to 2%, as mean ± SD (n=5). (**C**) The specificity of the m^6^A capture was tested by spiking poly(A) RNA from ESCs with exogenous RNAs before performing MeRIP-qPCR. Data are mean ± SD (n=3). Student ’s *t*-test \*\**P* < 0.01. (**D**) Examples of gene track views of MeRIP-seq, for mRNAs of pluripotency factors previously shown to be methylated in ESCs. (**E**) Motif analysis from MeRIP-seq peaks identifies several motifs corresponding to the consensus “DRACH” m^6^A motif (where D=A, G or U; H=A, C or U). (**F**) Distribution of differential m^6^A peaks, according to the type of structural element within the transcript. (**G**) Examples of gene track views of MeRIP-seq, for mRNAs with significant hypermethylation (*Tial1, Ptrf*) or hypomethylation (*Cenpt*) in pausing of ESCs.

**Fig. S4.**
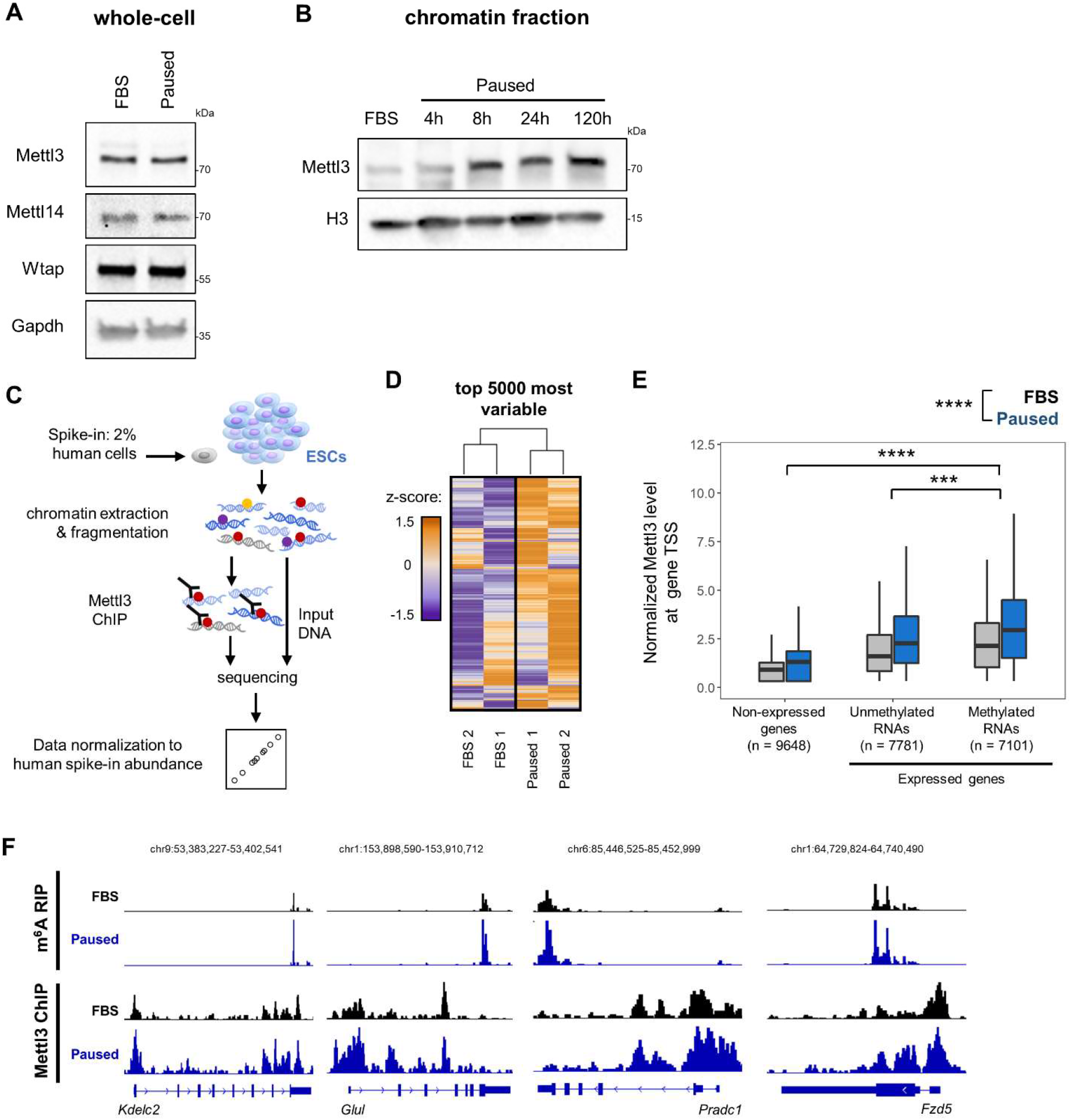
Mapping the chromatin distribution of Mettl3 in paused ESCs. (**A**) m^6^A machinery in control (FBS) and paused ESCs by western blot in whole cell extracts. (**B**) Increase of Mettl3 levels in chromatin extracts upon induction of paused pluripotency, measured by cell number-normalized (CNN) western blot. (**C**) Strategy for Mettl3 ChIP-seq in ESCs with CNN approach using human cell spiking. (**D**) Heatmap of the top 5000 most variable Mettl3 peaks across all samples, showing higher levels in paused ESCs. (**E**) Average levels of Mettl3 binding in the TSS of all genes, separated by expression and methylation status, in FBS and paused ESCs. Mettl3 binding is highest in the TSS of expressed genes with a methylated transcript, and in paused ESCs. Linear regression test \*\*\**P* < 0.001, \*\*\*\**P* < 0.0001. (**F**) Examples of gene track views showing increased levels (log_2_ fold-change > 0) of m^6^A and Mettl3.

**Fig. S5:**
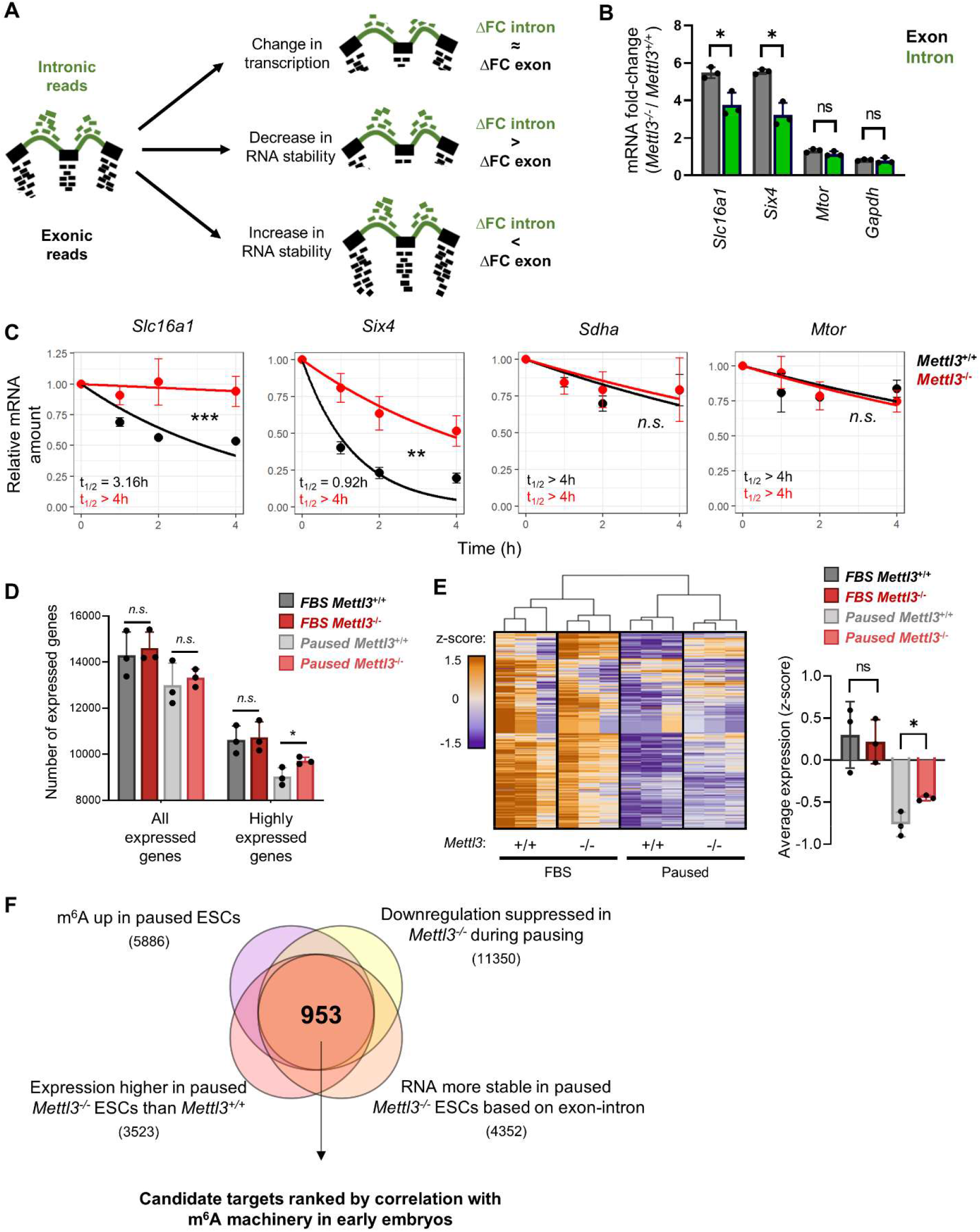
Exon-intron analysis captures Mettl3-dependent changes in RNA stability in paused ESCs. (**A**) Strategy for RNA stability analysis based on intronic and exonic reads. (**B**) Examples of genes with different (*Slc16a1, Six4*) and similar (*Mtor, Gapdh*) intronic and exonic mRNA fold-changes between *Mettl3*^*-/-*^ and *Mettl3*^*+/+*^ ESCs based on RNA-seq data. Student ’s *t*-tests \**P* < 0.05. (**C**) Validation of stability changes by actinomycin D stability assay (n=3). t_1/2_: half-life. Linear regression test \*\**P* < 0.01, \*\*\**P* < 0.001. (**D**) Quantification of the number of expressed genes in *Mettl3*^*+/+*^ and *Mettl3*^*-/-*^ ESCs based on intronic RNA-seq, in FBS and paused conditions. Expressed genes are further defined as having high expression (log_2_ normalized reads > 5). Data are mean ± SD (n=3). Student ’s *t*-tests \**P* < 0.05. (**E**) Heatmap of gene expression based on intronic reads for all genes expressed in *Mettl3*^*+/+*^ or *Mettl3*^*-/-*^ ESCs with average expression per sample (scored as median z-scores of all genes), showing defective hypotranscription in paused *Mettl3*^*-/-*^ ESCs. Student ’s *t*-tests \**P* < 0.05. (**F**) Identification of putative anti-pausing factors kept in check by m^6^A methylation and thereby destabilization of their transcript in paused pluripotency (see Methods for details).

**Fig. S6:**
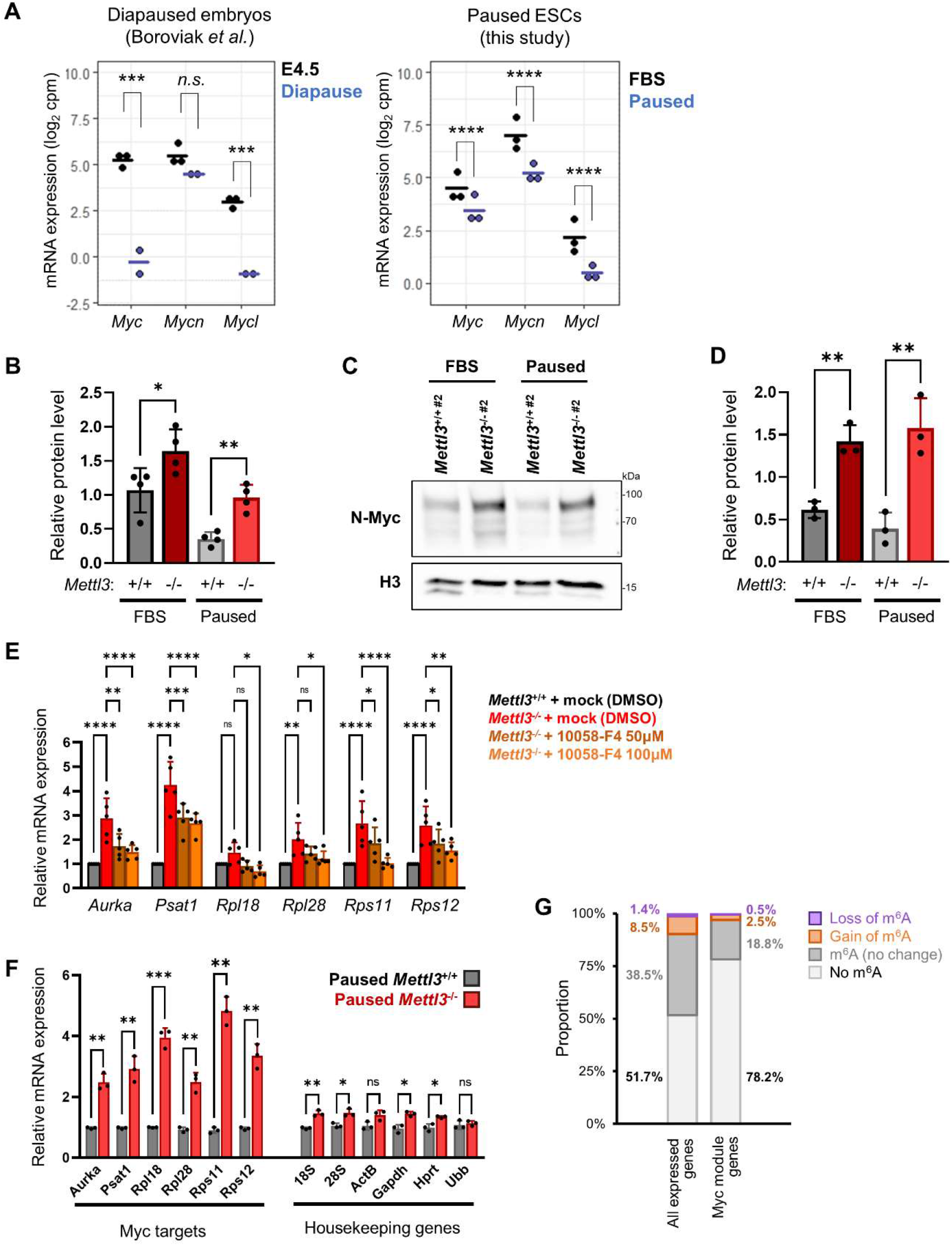
Regulation of Myc family members and downstream targets by Mettl3 in paused pluripotency. (**A**) Expression levels of the Myc factors in diapaused embryos (Boroviak *et al*.) and paused ESCs. Student ’s *t*-tests \*\*\**P* < 0.001, \*\*\*\**P* < 0.0001. (**B**) Quantification of N-Myc western blots, showing increased expression in *Mettl3*^*-/-*^ ESCs, as shown in Fig. 4B. Data are mean ± SD (n=4). Student ’s *t*-tests \**P* < 0.05, \*\**P* < 0.01. (**C**,**D**) Representative western blot and quantification of N-Myc protein levels in *Mettl3*^*-/- #2*^ ESCs. Data are mean ± SD (n=3). Student ’s *t*-tests \*\**P* < 0.01. (**E**) Blocking of Myc signaling partially restores the expression of Myc target genes in paused *Mettl3*^-/-^ ESCs. Data are mean ± SD (n=5). Two-way ANOVA with Dunnett ’s multiple comparison tests \**P* < 0.05, \*\**P* < 0.01, \*\*\**P* < 0.001, \*\*\*\**P* < 0.0001. (**F**) Nascent RNA capture by EU incorporation for Myc target genes and control housekeeping genes in paused *Mettl3*^*+/+*^ and *Mettl3*^-/-^ ESCs. Data are mean ± SD (n=3). Student ’s *t*-tests \**P* < 0.05, \*\**P* < 0.01, \*\*\**P* < 0.001. (**G**) The majority Myc module genes are not direct targets of m^6^A in paused pluripotency. The proportion in all genes expressed in ESCs is shown for comparison.

**Fig. S7.**
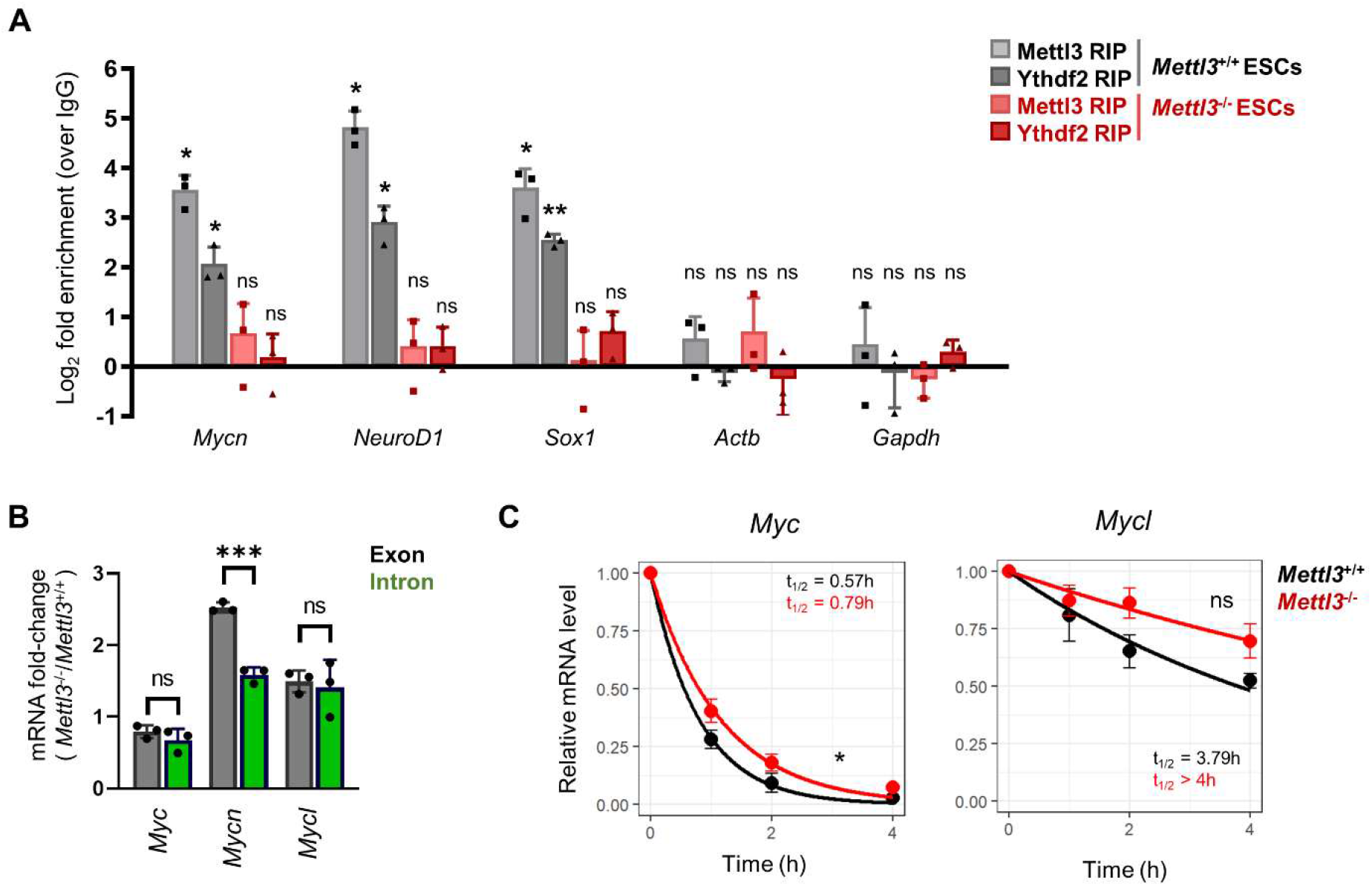
m_6_A-specific regulation of *Mycn* mRNA stability. (**A**) Mettl3 and Ythdf2 binding of the *Mycn* transcript, measured by RIP-qPCR. *NeuroD1* and *Sox2* were used as positive controls, and *Actb* and *Gapdh* were used as negative controls. Data are mean ± SD (n=3). Two-way ANOVA with Dunnett ’s multiple comparison tests \**P* < 0.05, \*\**P* < 0.01. (**B**) *Mycn* is the only Myc family member regulated at the RNA stability level by Mettl3, as evidenced by analysis of exonic and intronic mRNA fold-changes (*Mettl3*^*-/-*^/*Mettl3*^*+/+-*^ ESCs). Data are mean ± SD (n=3). Student ’s *t*-tests \*\*\**P* < 0.001. (**C**) Unlike *Mycn, Myc* and *Mycl* mRNAs display minimal changes in mRNA stability in *Mettl3*^*-/-*^ ESCs by actinomycin D assay. Data are mean ± SEM (n=3). Linear regression test \**P* < 0.05.

**Fig. S8.**
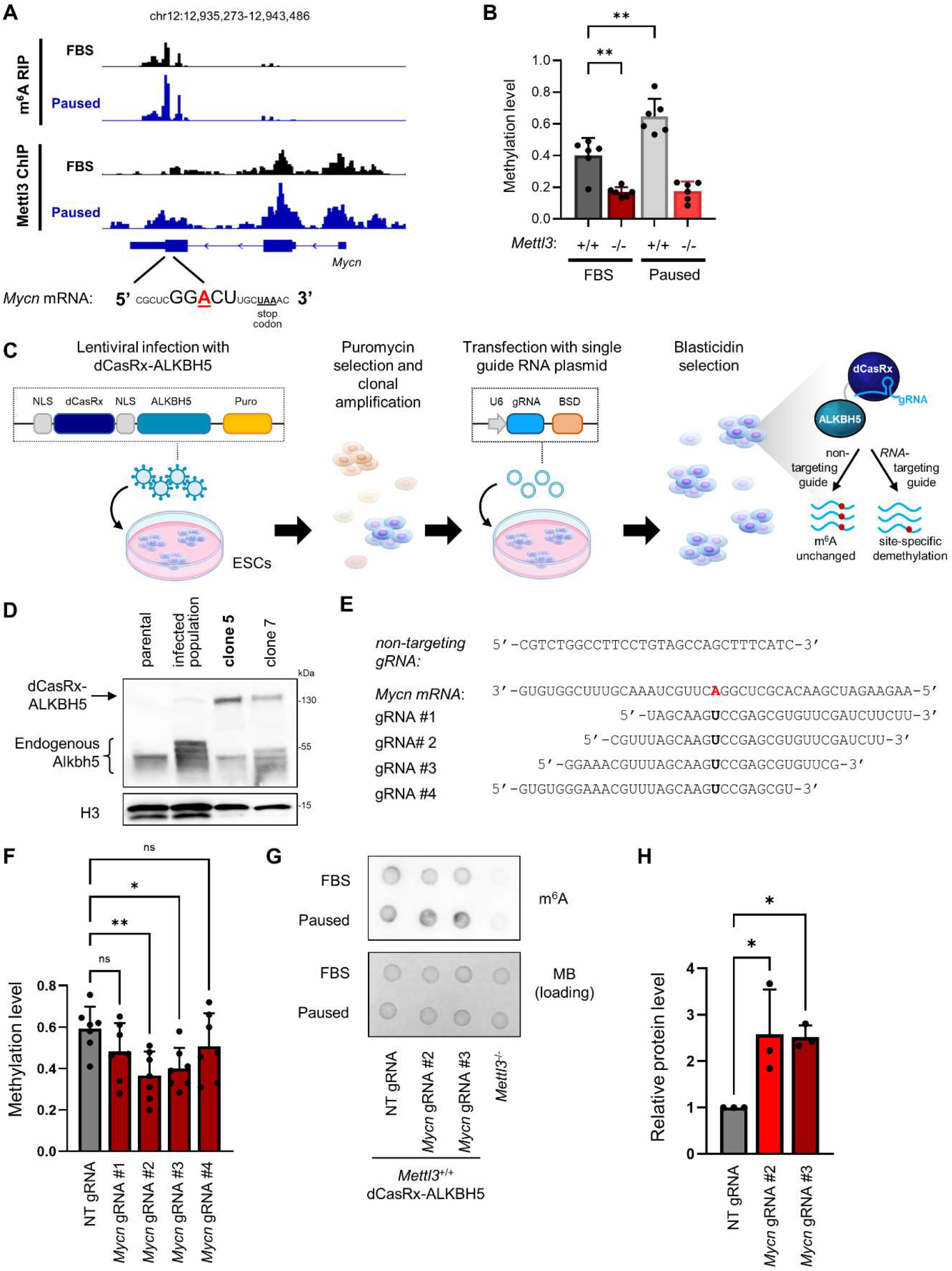
Targeted m_6_ A demethylation controls expression of *Mycn* in paused ESCs. (**A**) Gene track view of MeRIP-seq and Mettl3 ChIP-seq for *Mycn* mRNA. (**B**) Validation of m^6^A changes in *Mettl3*^*+/+*^ and *Mettl3*^*-/-*^ ESCs by m^6^A-qPCR. Data are mean ± SD (n=6). One-way ANOVA with Dunnett ’s multiple comparison tests \*\**P* < 0.01. (**C**) Model of lentiviral dCasRx epitranscriptomic editor with the m^6^A demethylase ALKBH5 and single guide RNA. (**D**) Validation of the expression of the dCasRx-ALKBH5 fusion by western blot in parental (non-infected) ESCs, infected ESCs, and 2 infected clones. Clone 5 was used for all experiments. (**E**) Guide RNAs (gRNAs) transfected for non-targeting control and *Mycn*-targeting conditions. (**F**). Guides #2 and #3 significantly reduce m^6^A in *Mycn* transcripts, as measured by m^6^A-qPCR. Data are mean ± SD (n=7). One-way ANOVA with Dunnett ’s multiple comparison tests \**P* < 0.05, \*\**P* < 0.01. (**G**) Dot blot showing that the global increase of m^6^A in paused ESCs is not affected by dCasRx-ALKBH5. MB: Methylene blue. (**H**) Quantification of N-Myc protein levels, showing increased expression with dCasRx-ALKBH5 targeting *Mycn* in paused ESCs. Data are mean ± SD (n=3). One-way ANOVA with Dunnett ’s multiple comparison tests \**P* < 0.05.

## Supplementary Tables

**Table S1. Differential expression analysis from RNA-seq data**.

Differential analysis of gene expression (exonic reads) upon pausing was performed separately for *Mettl3*^+/+^ and *Mettl3*^-/-^ ESCs using edgeR in R. The threshold for significant differential expression was adjusted *P* < 0.05 and fold-change > 1.5. Data were cell-number normalized. (Provided as a separate supplementary online Excel file)

**Table S2. Gene Set Enrichment Analysis using RNA-seq data**.

GSEA analysis was performed from RNA-seq data using the fGSEA package in R, pre-ranking genes by *t*-values from the differential analysis (paused/FBS). NES: normalized enrichment score. Gene set collections were downloaded from the Molecular Signatures Database (http://www.gsea-msigdb.org/gsea/msigdb/index.jsp).

(Provided as a separate supplementary online Excel file)

**Table S3. Differential methylation analysis from MeRIP-seq data**.

Differential analysis of m^6^A RNA methylation upon pausing was performed in wildtype ESCs using edgeR in R. The threshold for significant differential expression was adjusted *P* < 0.05 and fold-change > 1.5. Data were cell-number normalized.

(Provided as a separate supplementary online Excel file)

**Table S4. Differential expression analysis from RNA-seq data (intron)**.

Differential analysis of gene expression (intronic reads) upon pausing was performed separately for *Mettl3*^+/+^ and *Mettl3*^-/-^ ESCs using edgeR in R. The threshold for significant differential expression was adjusted *P* < 0.05 and fold-change > 1.5. Data were cell-number normalized.

(Provided as a separate supplementary online Excel file)

**Table S5. Top 100 ranked m**_**6**_**A anti-pausing candidates in paused pluripotency**.

Putative anti-pausing genes were ranked by their Spearman correlation coefficient (ρ) with an expression signature of the m^6^A machinery (see Methods for detail). The top 100 ranked genes are shown here.

(Provided as a separate supplementary online Excel file)

**Table S6.**
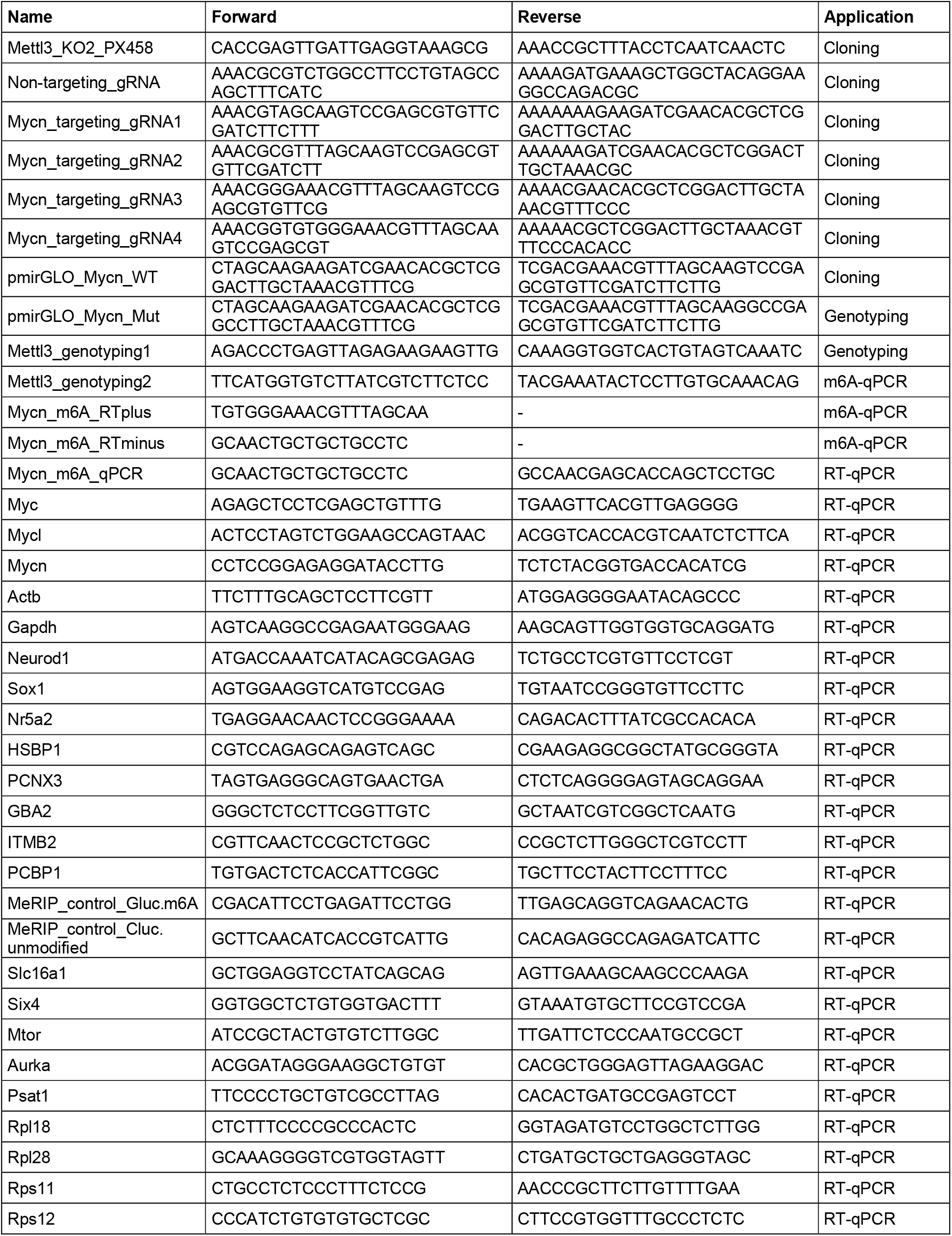
List of primers.

**Table S7.** List of antibodies.

| **Name** | **Source** | **Catalog number**<br>**(RRID if available)** | **Dilution or**<br>**amount** | **Application** |
| --- | --- | --- | --- | --- |
| Mettl3 | Abcam | ab195352<br>(AB_2721254) | WB 1:1000<br>IF: 1/200<br>RIP: 2µg | Western blot,<br>Immunofluorescence,<br>RIP |
| N-Myc | Cell Signaling<br>Technology | D1V2A | WB 1:1000<br>IF: 1/200 | Western blot,<br>Immunofluorescence |
| HRP-Goat Anti-Rabbit IgG (H+L) | Jackson Labs | 111-035-144<br>(AB_2307391) | 1:5000 | Western blot, dot<br>blot (secondary) |
| HRP-Goat Anti-Mouse IgG (H+L) | Jackson Labs | 115-035-062<br>(AB_2338504) | 1:5000 | Western blot<br>(secondary) |
| H3 | Abcam | ab1791 (AB_302613) | 1:2000 | Western blot |
| Mettl14 | Proteintech | 26158-1-AP<br>(AB_2800447) | 1:1000 | Western blot |
| Wtap | Proteintech | 60188-1-Ig<br>(AB_10859484) | 1:1000 | Western blot |
| Ythdf2 | Millipore Sigma | ABE542 | 2µg | RIP |
| Normal Rabbit IgG<br>Polyclonal | Millipore Sigma | 12-370 (AB_145841) | 2µg | RIP |
| Donkey anti-Rabbit<br>IgG (H+L) Highly<br>Cross-Adsorbed<br>Secondary Antibody,<br>Alexa Fluor™ 594 | Invitrogen | A-21207<br>(AB_141637) | 1:1000 | Immunofluorescence<br>(secondary) |
| m6A | Diagenode | C15410208 | 1:400 | Dot blot |
| Mettl3 | Proteintech | 15073-1-AP<br>(AB_2142033) | 5µg | ChIP |

## References

1. M. B. Renfree, J. C. Fenelon, The enigma of embryonic diapause. Development. 144, 3199–3210 (2017).

2. V. A. van der Weijden, A. Bulut-Karslioglu, Molecular Regulation of Paused Pluripotency in Early Mammalian Embryos and Stem Cells. Front. Cell Dev. Biol. 9, 2039 (2021).

3. J. C. Fenelon, M. B. Renfree, The history of the discovery of embryonic diapause in mammals. Biol. Reprod. 99, 242–251 (2018).

4. T. Boroviak, R. Loos, P. Lombard, J. Okahara, R. Behr, E. Sasaki, J. Nichols, A. Smith, P. Bertone, Lineage-Specific Profiling Delineates the Emergence and Progression of Naive Pluripotency in Mammalian Embryogenesis. Dev. Cell. 35, 366– 382 (2015).

5. A. Bulut-Karslioglu, S. Biechele, H. Jin, T. A. MacRae, M. Hejna, M. Gertsenstein, J. S. Song, M. Ramalho-Santos, Inhibition of mTOR induces a paused pluripotent state. Nature. 540, 119–123 (2016).

6. P. Wang, K. A. Doxtader, Y. Nam, Structural Basis for Cooperative Function of Mettl3 and Mettl14 Methyltransferases. Mol. Cell. 63, 306–317 (2016).

7. D. Dominissini, S. Moshitch-Moshkovitz, S. Schwartz, M. Salmon-Divon, L. Ungar, S. Osenberg, K. Cesarkas, J. Jacob-Hirsch, N. Amariglio, M. Kupiec, R. Sorek, G. Rechavi, Topology of the human and mouse m6A RNA methylomes revealed by m6A-seq. Nature. 485, 201–206 (2012).

8. P. J. Batista, B. Molinie, J. Wang, K. Qu, J. Zhang, L. Li, D. M. Bouley, E. Lujan, B. Haddad, K. Daneshvar, A. C. Carter, R. A. Flynn, C. Zhou, K. S. Lim, P. Dedon, M. Wernig, A. C. Mullen, Y. Xing, C. C. Giallourakis, H. Y. Chang, m6A RNA Modification Controls Cell Fate Transition in Mammalian Embryonic Stem Cells. Cell Stem Cell. 15, 707–719 (2014).

9. S. Geula, S. Moshitch-Moshkovitz, D. Dominissini, A. A. Mansour, N. Kol, M. Salmon-Divon, V. Hershkovitz, E. Peer, N. Mor, Y. S. Manor, M. S. Ben-Haim, E. Eyal, S. Yunger, Y. Pinto, D. A. Jaitin, S. Viukov, Y. Rais, V. Krupalnik, E. Chomsky, M. Zerbib, I. Maza, Y. Rechavi, R. Massarwa, S. Hanna, I. Amit, E. Y. Levanon, N. Amariglio, N. Stern-Ginossar, N. Novershtern, G. Rechavi, J. H. Hanna, Stem cells. m6A mRNA methylation facilitates resolution of naïve pluripotency toward differentiation. Science. 347, 1002–6 (2015).

10. Y. Wang, Y. Li, J. I. Toth, M. D. Petroski, Z. Zhang, J. C. Zhao, N6-methyladenosine modification destabilizes developmental regulators in embryonic stem cells. Nat. Cell Biol. 16, 191–8 (2014).

11. M. Percharde, A. Bulut-Karslioglu, M. Ramalho-Santos, Hypertranscription in Development, Stem Cells, and Regeneration. Dev. Cell (2017),, doi:10.1016/j.devcel.2016.11.010.

12. A. M. Hussein, Y. Wang, J. Mathieu, L. Margaretha, C. Song, D. C. Jones, C. Cavanaugh, J. W. Miklas, E. Mahen, M. R. Showalter, W. L. Ruzzo, O. Fiehn, C. B. Ware, C. A. Blau, H. Ruohola-Baker, Metabolic Control over mTOR-Dependent Diapause-like State. Dev. Cell. 52, 236–250 (2020).

13. R. Scognamiglio, N. Cabezas-Wallscheid, M. C. Thier, S. Altamura, A. Reyes, Á. M. Prendergast, D. Baumgärtner, L. S. Carnevalli, A. Atzberger, S. Haas, L. Von Paleske, T. Boroviak, P. Wörsdörfer, M. A. G. Essers, U. Kloz, R. N. Eisenman, F. Edenhofer, P. Bertone, W. Huber, F. Van Der Hoeven, A. Smith, A. Trumpp, Myc Depletion Induces a Pluripotent Dormant State Mimicking Diapause. Cell. 164, 668–680 (2016).

14. M. I. Sousa, B. Correia, A. S. Rodrigues, J. Ramalho-Santos, Metabolic characterization of a paused-like pluripotent state. Biochim. Biophys. acta. Gen. Subj. 1864 (2020), doi:10.1016/J.BBAGEN.2020.129612.

15. I. Barbieri, K. Tzelepis, L. Pandolfini, J. Shi, G. Millán-Zambrano, S. C. Robson, D. Aspris, V. Migliori, A. J. Bannister, N. Han, E. De Braekeleer, H. Ponstingl, A. Hendrick, C. R. Vakoc, G. S. Vassiliou, T. Kouzarides, Promoter-bound METTL3 maintains myeloid leukaemia by m6A-dependent translation control. Nature. 552, 126–131 (2017).

16. A. Bertero, S. Brown, P. Madrigal, A. Osnato, D. Ortmann, L. Yiangou, J. Kadiwala, N. C. Hubner, I. R. De Los Mozos, C. Sadée, A. S. Lenaerts, S. Nakanoh, R. Grandy, E. Farnell, J. Ule, H. G. Stunnenberg, S. Mendjan, L. Vallier, The SMAD2/3 interactome reveals that TGFβ controls m6A mRNA methylation in pluripotency. Nature. 555, 256–259 (2018).

17. W. Xu, J. Li, C. He, J. Wen, H. Ma, B. Rong, J. Diao, L. Wang, J. Wang, F. Wu, L. Tan, Y. G. Shi, Y. Shi, H. Shen, METTL3 regulates heterochromatin in mouse embryonic stem cells. Nature. 591, 317–321 (2021).

18. D. Gaidatzis, L. Burger, M. Florescu, M. B. Stadler, Analysis of intronic and exonic reads in RNA-seq data characterizes transcriptional and post-transcriptional regulation. Nat. Biotechnol. 33, 722–729 (2015).

19. T. R. Kress, A. Sabò, B. Amati, MYC: connecting selective transcriptional control to global RNA production. Nat. Rev. Cancer. 15, 593–607 (2015).

20. M. Percharde, P. Wong, M. Ramalho-Santos, Global Hypertranscription in the Mouse Embryonic Germline. Cell Rep. 19, 1987–1996 (2017).

21. I. A. Roundtree, M. E. Evans, T. Pan, C. He, Dynamic RNA Modifications in Gene Expression Regulation. Cell. 169, 1187–1200 (2017).

22. Z. Xia, M. Tang, J. Ma, H. Zhang, R. C. Gimple, B. C. Prager, H. Tang, C. Sun, F. Liu, P. Lin, Y. Mei, R. Du, J. N. Rich, Q. Xie, Epitranscriptomic editing of the RNA N6-methyladenosine modification by dCasRx conjugated methyltransferase and demethylase. Nucleic Acids Res. 49, 7361–7374 (2021).

23. J. Liu, X. Dou, C. Chen, C. Chen, C. Liu, M. Michelle Xu, S. Zhao, B. Shen, Y. Gao, D. Han, C. He, N 6-methyladenosine of chromosome-associated regulatory RNA regulates chromatin state and transcription. Science. 367, 580–586 (2020).

24. T. Chelmicki, E. Roger, A. Teissandier, M. Dura, L. Bonneville, S. Rucli, F. Dossin, C. Fouassier, S. Lameiras, D. Bourc‘his, m 6 A RNA methylation regulates the fate of endogenous retroviruses. Nature. 591, 312–316 (2021).

25. J. Wei, X. Yu, L. Yang, X. Liu, B. Gao, B. Huang, X. Dou, J. Liu, Z. Zou, X.-L. Cui, L.-S. Zhang, X. Zhao, Q. Liu, P. C. He, C. Sepich-Poore, N. Zhong, W. Liu, Y. Li, X. Kou, Y. Zhao, Y. Wu, X. Cheng, C. Chen, Y. An, X. Dong, H. Wang, Q. Shu, Z. Hao, T. Duan, Y.-Y. He, X. Li, S. Gao, Y. Gao, C. He, FTO mediates LINE1 m6A demethylation and chromatin regulation in mESCs and mouse development. Science. 376, 968–973 (2022).

26. M. Laplante, D. M. Sabatini, mTOR signaling in growth control and disease. Cell. 149, 274–293 (2012).

27. S. K. Rehman, J. Haynes, E. Collignon, K. R. Brown, Y. Wang, A. M. L. Nixon, J. P. Bruce, J. A. Wintersinger, A. Singh Mer, E. B. L. Lo, C. Leung, E. Lima-Fernandes, N. M. Pedley, F. Soares, S. McGibbon, H. H. He, A. Pollet, T. J. Pugh, B. Haibe-Kains, Q. Morris, M. Ramalho-Santos, S. Goyal, J. Moffat, C. A. O‘Brien, Colorectal Cancer Cells Enter a Diapause-like DTP State to Survive Chemotherapy. Cell. 184, 226-242.e21 (2021).

28. E. Dhimolea, R. de Matos Simoes, D. Kansara, A. Al‘Khafaji, J. Bouyssou, X. Weng, S. Sharma, J. Raja, P. Awate, R. Shirasaki, H. Tang, B. J. Glassner, Z. Liu, D. Gao, J. Bryan, S. Bender, J. Roth, M. Scheffer, R. Jeselsohn, N. S. Gray, I. Georgakoudi, F. Vazquez, A. Tsherniak, Y. Chen, A. Welm, C. Duy, A. Melnick, B. Bartholdy, M. Brown, A. C. Culhane, C. S. Mitsiades, An Embryonic Diapause-like Adaptation with Suppressed Myc Activity Enables Tumor Treatment Persistence. Cancer Cell. 39, 240-256.e11 (2021).

29. F. A. Ran, P. D. Hsu, J. Wright, V. Agarwala, D. A. Scott, F. Zhang, Genome engineering using the CRISPR-Cas9 system. Nat. Protoc. 8, 2281–2308 (2013).

30. T. A. Macrae, M. Ramalho-Santos, The deubiquitinase Usp9x regulates PRC2-mediated chromatin reprogramming during mouse development. Nat. Commun. 12, 1– 15 (2021).

31. R. Ross, X. Cao, N. Yu, P. A. Limbach, Sequence mapping of transfer RNA chemical modifications by liquid chromatography tandem mass spectrometry. Methods. 107, 73–78 (2016).

32. B. Delatte, F. Wang, L. V. Ngoc, E. Collignon, E. Bonvin, R. Deplus, E. Calonne, B. Hassabi, P. Putmans, S. Awe, C. Wetzel, J. Kreher, R. Soin, C. Creppe, P. A. Limbach, C. Gueydan, V. Kruys, A. Brehm, S. Minakhina, M. Defrance, R. Steward, F. Fuks, Transcriptome-wide distribution and function of RNA hydroxymethylcytosine. Science. 351, 282–285 (2016).

33. S. P. DiTroia, M. Percharde, M. Guerquin, E. Wall, E. Collignon, K. T. Ebata, K. Mesh, S. Mahesula, M. Agathocleous, D. J. Laird, G. Livera, M. Ramalho-Santos, Maternal Vitamin C regulates reprogramming of DNA methylation and germline development. Nature. 573, 271–275 (2019).

34. X. Wang, B. S. Zhao, I. A. Roundtree, Z. Lu, D. Han, H. Ma, X. Weng, K. Chen, H. Shi, C. He, N6-methyladenosine modulates messenger RNA translation efficiency. Cell. 161, 1388–1399 (2015).

35. J. Jeschke, E. Collignon, C. Al Wardi, M. Krayem, M. Bizet, Y. Jia, S. Garaud, Z. Wimana, E. Calonne, B. Hassabi, R. Morandini, R. Deplus, P. Putmans, G. Dube, N. K. Singh, A. Koch, K. Shostak, L. Rizzotto, R. L. Ross, C. Desmedt, Y. Bareche, F. Rothé, J. Lehmann-Che, M. Duterque-Coquillaud, X. Leroy, G. Menschaert, L. Teixeira, M. Guo, P. A. Limbach, P. Close, A. Chariot, E. Leucci, G. Ghanem, B. F. Yuan, K. Willard-Gallo, C. Sotiriou, J. C. Marine, F. Fuks, Downregulation of the FTO m6A RNA demethylase promotes EMT-mediated progression of epithelial tumors and sensitivity to Wnt inhibitors. Nat. Cancer. 2, 611–628 (2021).

36. Y. Tang, K. Chen, B. Song, J. Ma, X. Wu, Q. Xu, Z. Wei, J. Su, G. Liu, R. Rong, Z. Lu, J. P. D. Magalhães, D. J. Rigden, J. Meng, m6A-Atlas: a comprehensive knowledgebase for unraveling the N6-methyladenosine (m6A) epitranscriptome. Nucleic Acids Res. 49, D134–D143 (2021).

37. A. Castellanos-Rubio, I. Santin, A. Olazagoitia-Garmendia, I. Romero-Garmendia, A. Jauregi-Miguel, M. Legarda, J. R. Bilbao, A novel RT-QPCR-based assay for the relative quantification of residue specific m6A RNA methylation. Sci. Rep. 9, 1–7 (2019).

38. A. Olazagoitia-Garmendia, A. Castellanos-Rubio, Relative Quantification of Residue-Specific m6A RNA Methylation Using m6A-RT-QPCR. Methods Mol. Biol. 2298, 185–195 (2021).

39. H. B. Sadowski, M. Z. Gilman, Cell-free activation of a DNA-binding protein by epidermal growth factor. Nature. 362, 79–83 (1993).

40. J. Méndez, X. H. Zou-Yang, S. Y. Kim, M. Hidaka, W. P. Tansey, B. Stillman, Human origin recognition complex large subunit is degraded by ubiquitin-mediated proteolysis after initiation of DNA replication. Mol. Cell. 9, 481–491 (2002).

